# *AlzGenPred*: A CatBoost based method using network features to classify the Alzheimer’s Disease associated genes from the high throughput sequencing data

**DOI:** 10.1101/2023.11.04.565636

**Authors:** Rohit Shukla, Tiratha Raj Singh

## Abstract

**Background and Objective:** AD is a progressive neurodegenerative disorder characterized by memory loss. Due to the advancement in next-generation sequencing technologies, an enormous amount of AD-associated genomics data is available. However, the information about the involvement of these genes in AD association is still a research topic because all these algorithms are based on statistical techniques. Therefore, AlzGenPred is developed to identify the AD-associated genes from a large set of data.

**Methods:** To develop the AlzGenPred, we have compiled a benchmark dataset consisting of 1086 AD and non-AD genes and used them as positive and negative datasets. We have generated several features including the fused features and evaluated them through machine learning methods. Then hyperparameter tuning approach was also applied and the final model was selected. The proposed method was validated by using the AlzGene and transcriptomics datasets and proposed as a standalone tool.

**Results:** Total 13504 features belonging to eight different encoding schemes of these sequences were generated and evaluated by using 16 ML algorithms. It reveals that network-based features can classify AD genes while sequence-based features are not able to classify them. Then we generated 24 different fused features (6020 D) using sequence-based features and fed them into a two-step lightGBM-based recursive feature selection method. It increased up to 5-7% accuracy. After that selected eight fused features with CKSAAP were used for the hyperparameter tuning. They showed <70% accuracy. Therefore, network-based features were used to generate the CatBoost-based ML method called AlzGenPred with 96.55% accuracy and 98.99% AUROC. The developed method is tested on the AlzGene dataset where it showed 96.43% accuracy. Then the model is validated using the transcriptomics dataset also.

**Conclusion:** The validation of AlzGenPred using the AlzGene dataset and transcriptomics dataset obtained from Human, mouse, and ES-derived neural cells revealed that it can classify the omics data and can sort the AD-associated genes. These predicted genes can be directly used in the wet lab for further testing which will reduce labor cost and time expenses. The AlzGenPred is developed as a standalone package and is available for users at https://www.bioinfoindia.org/alzgenpred/ and https://github.com/shuklarohit815/AlzGenPred.

## 1. Introduction

Alzheimer’s disease (AD) is the most common cause of dementia characterized by cognitive impairment and memory loss [1]. AD is ranked the 6^th^ leading cause of death in the USA. It is estimated that approximately 50 million people will suffer from dementia in 2050 and 60-70% will suffer from AD. Almost 5.7 million Americans are already suffering from dementia. Currently, AD patient requires $232 billion expense for the take care, which is a great factor to increase the risk of negative mental and physical health and emotional anxiety issues to caregivers. Although, if AD can be characterized in its early stage, it can reduce medical expenses cost of $7.9 trillion and reduce care expenses also [2]. Currently, AD is mainly diagnosed by brain imaging, neurocognitive tests, and cerebrospinal fluid (CSF) assays [3,4]. The deposition level of neurofibrillary tangles (NFTs) and amyloid-beta plaques (Aβ) with significant synapse loss is the major parameter to assess the brain pathology of AD patients. Current diagnostic methods have included the level of total tau protein and hyperphosphorylated tau (p-tau) with the amyloid-β_1-42_ (Aβ_1_-_42_) in cerebrospinal fluid (CSF) [5–7]. Both CSF biomarkers are also used for research purposes but these methods are expensive [8,9]. Further, the specificity and sensitivity of these CSF biomarkers (Aβ_42_ and p-tau) have raised serious concerns about their clinical implications [5,6,10]. The specificity for CSF Aβ_42_ ranges from 0.44 to 0.89 and sensitivity ranges from 0.69 to 0.81 [11]. Although, generally AD is diagnosed in late stages and if we can early identify AD before the major changes in the brain or brain damage then AD patients can get more benefit from the medication and care. Hence biomarker identification which can early diagnose AD is a very important task because it can improve the diagnosis and can increase the treatment options at early stage without major AD progression [12].

Several genetic linkage studies, population surveys, and genome-wide association studies (GWAS) have been carried out to predict the AD-associated genes and their genetic mutations which can modify the gene expression pattern in the brain. Mutations in the genes such as amyloid precursor protein (APP), Presenilin-1 (PSEN1), Presenilin-2 (PSEN2), and Apolipoprotein E (ApoE) are the strongest risk factors for AD [13]. Various other genes such as Triggering receptor expressed on myeloid cells 2 (TREM2), sortilin-related receptor: L, phosphatidylinositol binding clatherin assembly protein 1, bone marrow stromal cell antigen 1, leucine-rich repeat kinase 2, complement receptor 1, and some genes showed a strong association with AD [14]. A lot of other genes have also required testing through traditional methods such as linkage studies and GWAS in populations which is time-consuming and requires a lot of effort, hence it is urgently required to develop some new method that can prioritize the potential genes and can reduce the size of genes generated from genomics techniques for lab testing [15]. Scientists are discovering some alternate approaches such as genomics, proteomics, machine learning, and other computational approaches which can decrease the number of genes for the experimental analysis. Recently some studies proposed methods for cancer gene identification [16–18] and similar approaches could help in devising new methods for AD and other neurological conditions.

The known AD genes to date, do not cover a significant portion of the human genome hence still an innumerable number of AD genes remain to be discovered. Although many AD genes are discovered, a lot of disease genes still need to reveal. As we know that no cure exists for AD hence identification of potential genes that can act as a biomarker and therapeutic targets will increase the progress of AD drug discovery. The network-based methods are also widely used to find the key genes from a large dataset that have an association with the disease. Jamal *et al.* used machine learning techniques on various features including protein-protein interaction (PPI) based features and proposed the potential genes for AD [19]. A recent study identified the infectious disease-associated host genes from the sequence and PPI-based features and prioritize the important genes [20]. The network-based approaches were also used to prioritize the disease-associated genes for several diseases [21,22].

We have used 16 machine learning algorithms to classify the AD and non-AD genes by using the sequence-based and PPI-based features. We have seen that CatBoost is performing well with the network-based features. We have also assessed the prediction efficiency of **AlzGenPred** on various independent datasets including gene expression data. Our method is predicting well on all these datasets therefore we have concluded that it can predict the AD-associated genes for any relevant blind dataset.

## 2. Material and Methods

We have depicted the complete design and evaluation process of the AlzGenPred method in **Figure 1**. The method comprises nine major steps: Dataset retrieval and pre-processing, feature extraction, feature selection, baseline model construction, feature fusion, hyperparameter tuning, best model selection, and implementation and last is method validation using external datasets validation.

**Figure 1.**
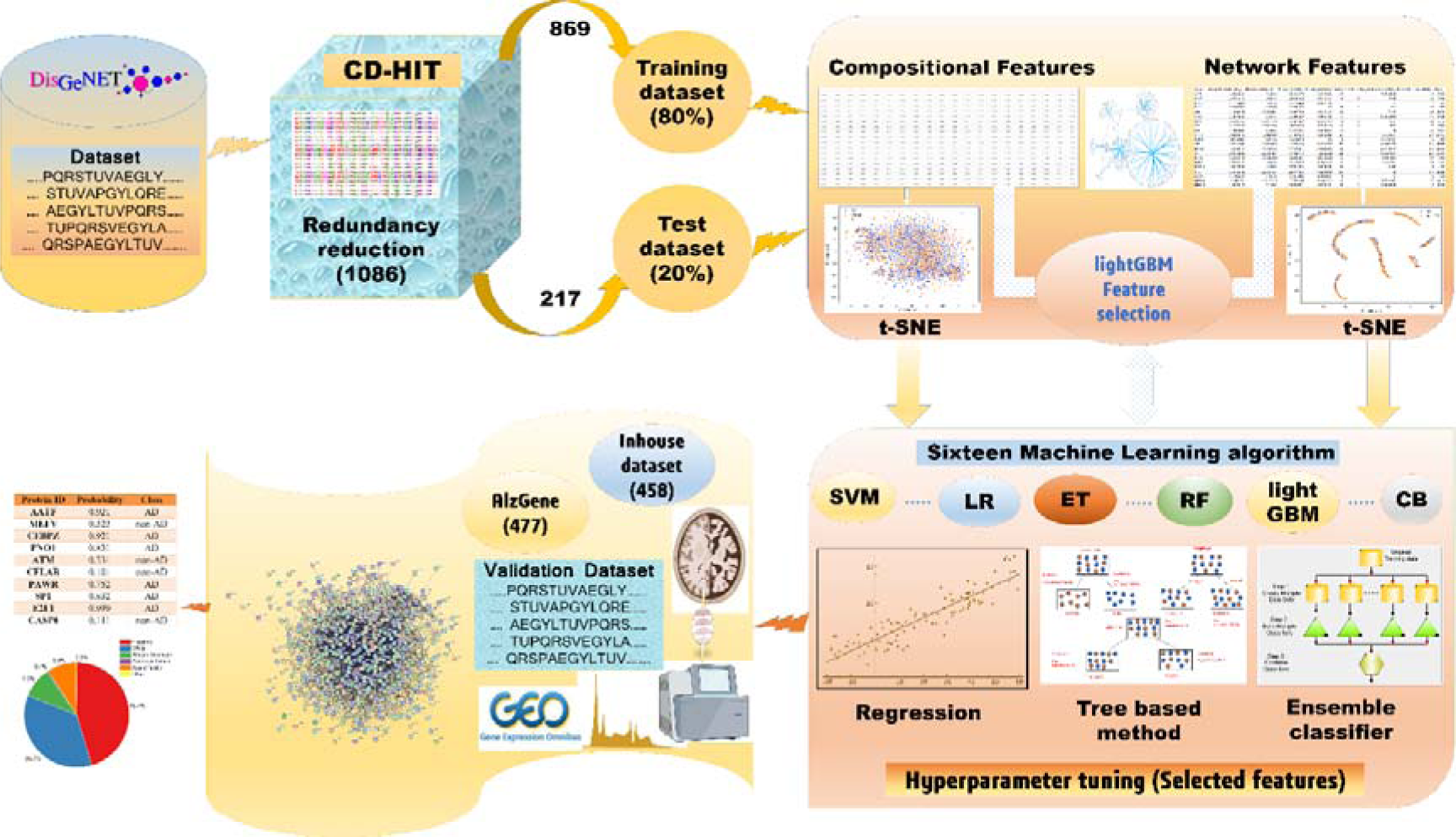
The comprehensive methodology of the work with all steps and their respective details.

### 2.1 Dataset collection and pre-processing

We have collected the AD-associated genes from the DisGeNET database [23]. It has expert-curated and disease-associated genes derived from text mining through various kinds of literature and public repositories. The database consists of information from various public databases such as Comparative Toxicogenomics Database (CTD) [24], GWAS Catalog [25], ClinVar [26], UniProtKB [27], Rat Genome Database (RGD) [28], Orphanet [29], Genetic Association Database (GAD) [30], BeFree data [31,32]. Mouse Genome Database (MGD) [33] and Literature Human Gene Derived Network (LHGDN) [34]. The term Alzheimer’s Disease (DisGeNET ID: C0002395) was used for the search in the DisGeNET database. It showed 3,397 genes associated with AD. We have only selected genes that showed the Evidence score (EI) (Gene disease association) >=1 and research publication >= 2. The EI score is calculated by the number of supporting literatures divided by the total number of literatures which also includes the articles which proved that these genes do not have any relation with this disease. The EI >= 1 describes that all the articles are showing the association of the gene with the disease while EI <= 1 represents that some literatures do not support the role of this gene in the disease. By filtering these criteria, we have selected 1270 AD genes. The genes were mapped to UniProt database and then 1270 sequences were retrieved from the UniProt. The homologous sequence should be removed to avoid the over fitting before training therefore, we have used a popular program CD-HIT [35] to remove the homologous AD sequences to cluster with the threshold of 40% sequence identity. If the two AD genes share >40% sequence identity then CD-HIT will keep only one representative sequence. On the basis of this criteria, we have found **1086** genes which does not have >40% identity between each other. This dataset is used as a positive dataset in this study.

We retrieved other neurodegenerative disease datasets from the DisGeNET and used them as a negative dataset in this study. We have selected several diseases such as Dementia (DisGeNET ID: C0497327), Huntington’s Disease (DisGeNET ID: C0020179), Parkinson’s Disease (DisGeNET ID: C0030567), and Pick’s Disease of the Brain (DisGeNET ID: C0236642). We merged the genes of all the diseases and removed the duplicates and non-coding genes etc. Hence, we found 2281 genes, and these selected genes were employed on the CD-HIT server to remove the homologous sequence. Based on 40% sequence identity we have found 1929 genes. We have taken dementia and other close diseases with the AD so they can share the same genes. Thus, we compared the negative dataset (1929 genes) with the positive dataset (1086 genes) and removed the identical sequence from the negative dataset. After the elimination of the common sequence, we found 1350 novel genes. We have used a balanced dataset for training the model hence we have randomly selected 1086 genes and treated them as a negative dataset.

Finally, the positive and negative datasets were divided into training (80%) and test (20%) to train the ML algorithms.

### 2.2 Feature encoding schemes

We have used eight different feature encoding algorithms from several perspectives to capture the key information of AD genes. The encoding schemes contain compositional-based feature information, autocorrelation-based features, conjoint triad and network topological features, etc. The description of the features with their dimensions is given in **Table 1**. To calculate the sequence-based features we have used the iFeature [36] toolkit based on Python 3.0 or above. We have extracted the sequence-based features based on the seven groups. The seven encoding schemes are briefly described below.

**Table 1.**
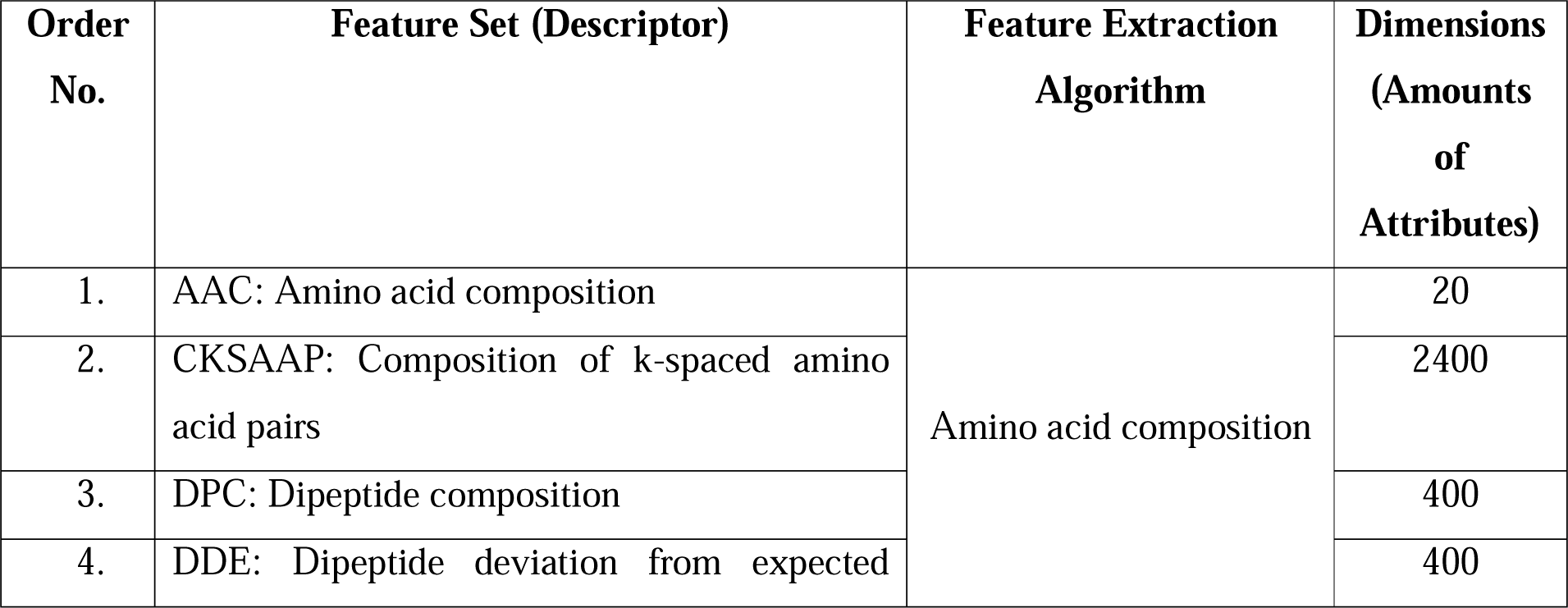
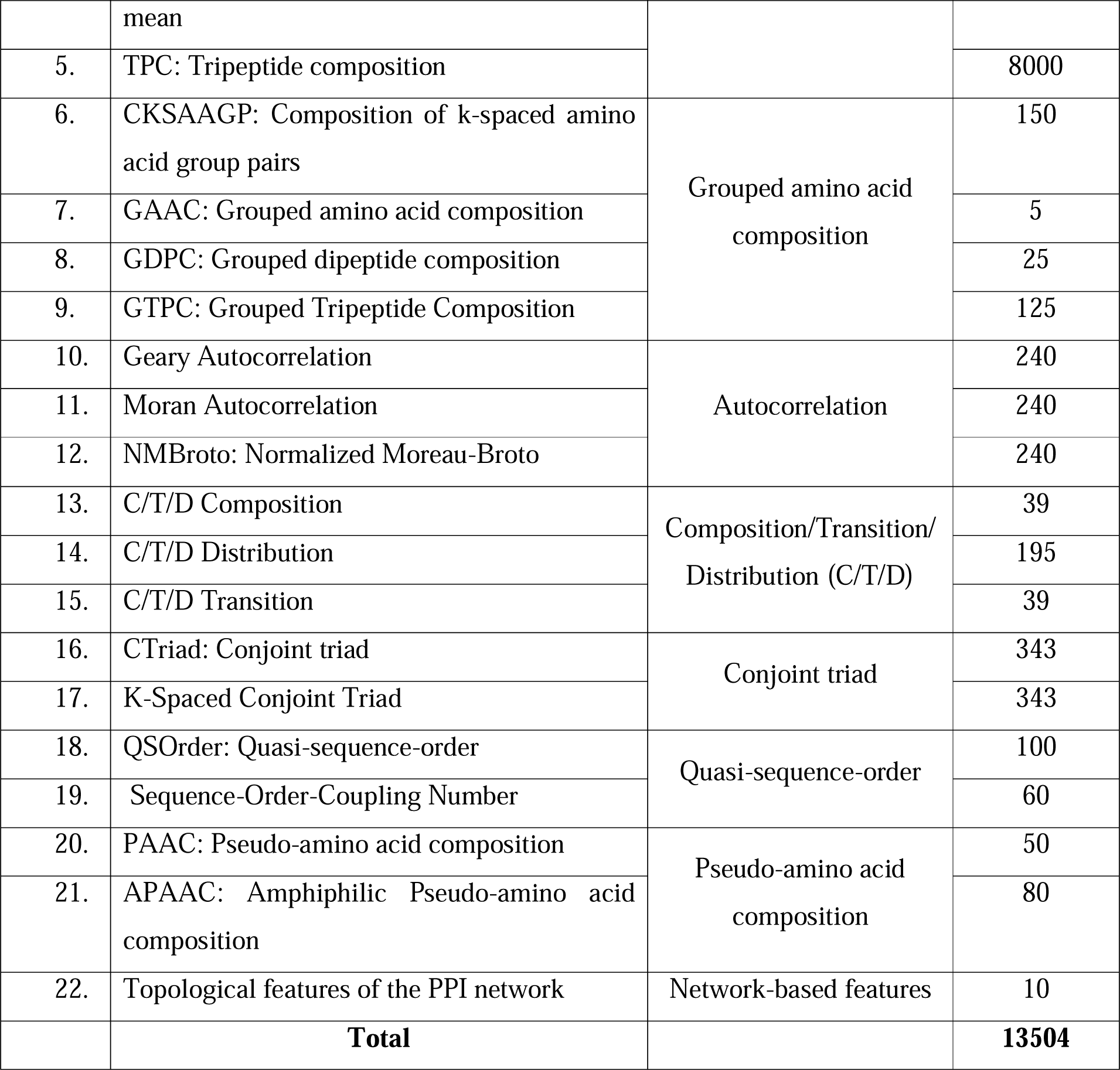
A summary of the selected 22 features.

#### 2.2.1 Amino acid composition

The Amino acid composition (AAC) is calculated by the normalization of each residue with the total number of residues present in the protein [37]. In this descriptor group, we have not just encoded each amino acid frequency in the AD genes to calculate the AAC while we have also calculated dipeptide composition (DPC), k-spaced amino acid pairs (CKSAAP), dipeptide deviation from the expected mean (DDE) and tripeptide composition (TPC) subsequently.

#### 2.2.2 Grouped amino acid composition

In this descriptor group, the features are categorized into five classes based on their physicochemical properties such as aliphatic, aromatic, positive charge amino acid, negative charge amino acid, and neutral amino acid [36]. We have not only calculated the GAAC but we have also calculated Grouped dipeptide composition (GDPC) and Composition of k-spaced amino acid group pairs (CKSAAGP). These features can provide a detailed result related to the charge, hydrophobicity, etc. for the AD genes.

#### 2.2.3 Autocorrelation

The Autocorrelation based features depend on the basis of the distribution of amino acid properties along the sequence and introduced by Moreau and Broto [38]. This descriptor group consists of three feature encoding schemes Moran, Geary, and Normalized Moreau-Broto (NMBroto).

#### 2.2.4 Composition**/**Transition**/**Distribution (C**/**T**/**D)

The amino acid distribution pattern of the AD proteins based on the specific physicochemical or structural property is represented by this descriptor group [39,40]. The features of this group are calculated by using the seven types of physical properties such as normalized Van der Waals volume, hydrophobicity, polarizability, charge, solvent accessibility, polarity, and secondary structures.

#### 2.2.5 Conjoint triad

By relating any three continuous amino acids as a distinct unit, this feature extractor may be used to investigate the properties of each amino acid in protein sequences and its vicinal amino acids [38].

#### 2.2.6 Quasi-sequence-order

Based on the augmented covariant discriminant technique, this promising descriptor can pass through the great difficulty of peptide/protein sequences (permutations and combinations). It also allows us to use diverse protein properties to attain improved prediction quality [41].

#### 2.2.7 Pseudo-amino acid composition

Chou has originally introduced the pseudo amino acid composition (PseAAC) descriptor group in 2001 to get the key features for improving the prediction of subcellular localization and membrane proteins [42]. This method also characterizes the protein by using an amino-acid frequency matrix of the protein sequence as compared to the original AAC method without significant sequential homology to other proteins.

#### 2.2.8 Network-based features

The Protein-Protein interaction (PPI) network of AD and non-AD genes was computed using the STRING database. All the selected 1086 genes were submitted to the STRING server and then the PPI was retrieved. After that, the retrieved PPI network was imported to Cytoscape [43] and then 10 topological features of each protein were calculated using the NetworkAnalyzer plugin [44]. These topological properties are AverageShortestPathLength, BetweennessCentrality, ClosenessCentrality, ClusteringCoefficient, Degree, Eccentricity, NeighborhoodConnectivity, NumberOfUndirectedEdges, Stress, and TopologicalCoefficient. The topological features were previously also used for the classification of AD and infectious diseases [19,20].

### 2.3 Two-step feature selection approach

Before feature selection, we considered all the features and fed them into 16 ML algorithms (CatBoost, Extra Trees, Linear Discriminant Analysis, Gradient Boosting Classifier, Bagging Classifier, Stochastic Gradient Descent Classifier, lightGBM, XGBoost, AdaBoost, Gaussian Naive Bayes, Multilayer perceptron, Decision Tree, Random Forest, Support Vector Machine, Logistic Regression and K-Nearest Neighbours) to generate the potential prediction models using 10-fold cross-validation. To identify the important feature set (optimal features) from the original features, the two-step feature selection approach is a systematic way and used in various studies [45–47]. In the first step, we ranked the original features by wrapping those features in an ML-based algorithm. To select the optimal features from biological data, gradient boosting methods such as Extreme Gradient Boosting (XgBoost) [48], etc. were used widely [49–51]. The recursive feature elimination method removes one feature and then checks the performance of the remaining feature set using a defined ML algorithm and then on the basis of correctly classified instance (accuracy) it proposes an optimal set of features. As these methods are very time-consuming and the speed depends on the selected ML algorithm. We have 13504 features from various feature schemes hence we have used lightGBM which is faster than XgBoost and takes less time for feature selection. The lightGBM ranks the features on the basis of their importance score and then gives the optimal subset of features. The top-ranked features predicted by the lightGBM were used in the second step. In the second step, we have inputted the selected feature subsets in 16 machine learning algorithms using 10-fold cross-validation. The machine learning models predicted from different algorithms and diverse sets of features were compared and then the best model for each feature set was selected based on accuracy. The best feature set was used for further study which showed the highest accuracy from one or more ML methods.

### 2.4 Feature fusion and feature selection

The feature fusion approach is a widely used and quite popular method in computational biology [52]. As we have already selected the best features from each encoding hence, we have only considered the selected features instead of the original features and made various combinations. We selected those feature sets which showed >=52% accuracy in the lightGBM feature selection method. Briefly, 8 encoding representation vectors (*E*) of AAC, DPC, DDE, GTPC, Geary, CTDD, PAAC, and CKSAAP were concatenated and 24 new features were created. The fused feature sets were again employed for the feature selection process and their evaluation using 16 ML algorithms.

### 2.5 Topological feature selection

Based on the above feature selection and model evaluations, we found that topological parameters were outperforming while sequence-based features are worst performing. We observed, that tree and gradient-boosting-based methods are performing best thus we only selected the topological features (10D) and selected two tree-based (Random Forest and Extra trees) and three gradients boosting based methods (XGBoost, lightGBM, and CatBoost) for further analysis. We have used the wrapper-based recursive feature elimination method for feature selection of topological features and then we calculated the accuracy of the selected optimal features by using these five methods. Three methods were selected with an optimal feature set (4D) for hyperparameter tuning which was the best performing smallest subset of features.

### 2.6 Hyperparameter optimization, model selection, and evaluation metrics

We have selected three algorithms (lightGBM, CatBoost, and ExtraTrees) with 4 feature subsets for network-based features. Eight fused features were also selected with CKSAAP (168D) for sequence-based features and used for hyperparameter tuning using eight ML algorithms (AdaBoost, CatBoost, ExtraTrees, Gradient Boosting, lightGBM, XGBoost, Linear Discriminant Analysis, and Multilayer perceptron). These algorithms have several parameters which can be optimized. We made a dictionary of parameters shown in **Supplementary Table S1** and supplied it to these eight classifiers and trained the model with 10-fold cross-validation. The GridSearchCV is a Scikit-learn package based on Python that is used to tune the parameter which can predict the best model from thousands of generated models.

The hyperparameter tuning result showed that CatBoost with 4D feature size is performing best as compared to other methods. Thus, the final model is selected and widely used metrics (accuracy (Acc), specificity (Sp), sensitivity (Sn), Precision, and Matthew’s correlation coefficient (MCC)) were used to evaluate the best model performance.

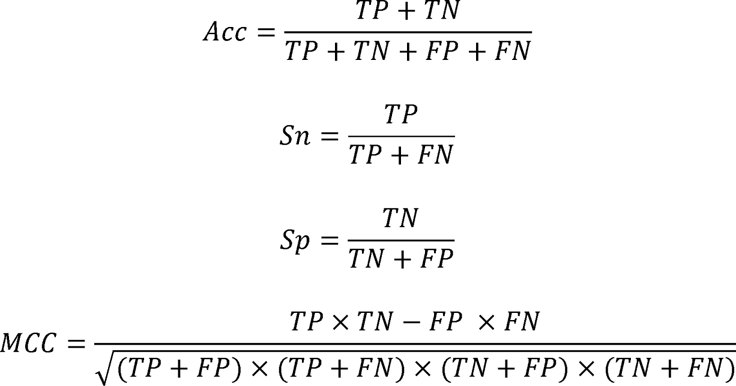

where TP, FP, TN, and FN represent the number of true positives, false positives, true negatives, and false negatives, respectively.

### 2.7 Method validation

The proposed AlzGenPred method was validated by using a diverse set of data. Firstly, we have taken the AlzGene dataset [53] (http://www.alzgene.org/) and their features were calculated and inputted into the method for the prediction. After that, we used the manually curated database for the model validation. Additionally, we have used the transcriptomics data for model validation. The Raw RNA-seq data for three datasets were retrieved from the European Nucleotide Archive (ENA) database [54]. These three datasets were generated from different samples but all belong to two conditions (AD *vs.* healthy control). The detail about the dataset is shown in **Table 2**. The quality of raw data was checked by using the FastQC toolkit (https://www.bioinformatics.babraham.ac.uk/projects/fastqc/). The low-quality reads and adapters were trimmed by using the TrimGalore tool (https://www.bioinformatics.babraham.ac.uk/projects/trim_galore/). The UCSC genome index file was used for the mapping. The trimmed reads were mapped to the respective genome (Human and Mouse) using the HiSat2 v2.1.0 [55]. The count files were generated by using the HTSeq count tool using the UCSC transcriptome file [56]. After that, we removed the lowly expressed genes from all three datasets.

**Table 2.**
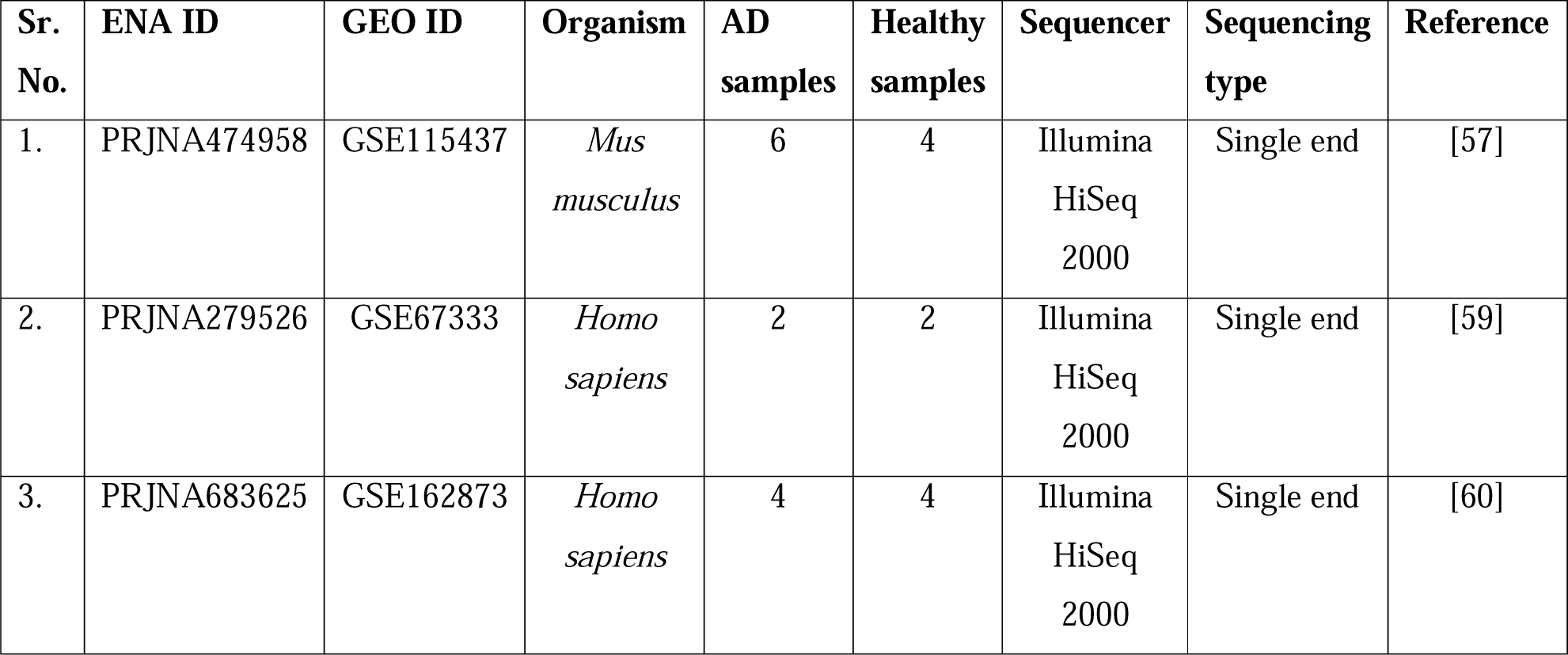
Summary of the raw RNAseq data used for the model validation.

| Sr. No. | ENA ID | GEO ID | Organism | AD samples | Healthy samples | Sequencer | Sequencing type | Reference |
| --- | --- | --- | --- | --- | --- | --- | --- | --- |
| 1. | PRJNA474958 | GSE115437 | *Mus musculus* | 6 | 4 | Illumina HiSeq 2000 | Single end | [57] |
| 2. | PRJNA279526 | GSE67333 | *Homo sapiens* | 2 | 2 | Illumina HiSeq 2000 | Single end | [59] |
| 3. | PRJNA683625 | GSE162873 | *Homo sapiens* | 4 | 4 | Illumina HiSeq 2000 | Single end | [60] |

For the first dataset [57] (GSE115437) generated from *Mus musculus* we have removed the reads which showed 0.5 CPM in at least one library using the edgeR tool [58]. The genes with log2fc > 1.0 and adjusted p-value or FDR (padj) < 0.05 were considered upregulated and overexpressed for the AD while genes with log2fc < −1.0 and padj < 0.05 were considered downregulated or under-expressed in AD patients. The samples of the second dataset [59] (GSE67333) were taken from Humans for AD and healthy control. We have removed the samples which showed the higher batch effect. Then the same pre-processing steps were used and the lowly expressed genes (2.0 CPM in at least two libraries) were removed from the count files. The differentially expressed genes (DEGs) were identified using a cut-off of padj < 0.05 and log2fc > 2.0 for overexpressed genes while padj < 0.05 and log2fc > −2.0 were used for the under-expressed genes. The third sample [60] (GSE162873) were generated from the ES-derived neural cell for the healthy and AD conditions. We have removed the lowly expressed genes which showed 1.0 CPM in at least 2 libraries. The DEGs were identified using a cut-off of padj < 0.05 and log2fc >2.0 for overexpressed genes while padj < 0.05 and log2fc >-2.0 were used for the underexpressed genes. After that, we did the gene set enrichment analysis and we observed that most of the pathways are related to AD. Then we utilized the overexpressed and underexpressed genes for the AD model validation. These genes were inputted to the STRING database (https://string-db.org/) and then PPI networks were generated. The generated map was fed to the NetAnalyzer plugin using the Cytoscape interface and then the topological features were generated and evaluated using the AlzGenPred tool to classify the AD *vs.* non-AD genes.

## 3. Results

### 3.1 Feature behavior analysis

Positive and negative datasets were taken from the neurodegenerative disease hence these may share a common pattern that will cause the problem in the model training. Therefore to better understand the feature space distribution we have used the *t*-distributed stochastic neighborhood embedding (t-SNE) [61] method to transform all the features in two-dimensional space to get a clear picture of the feature distribution. All 21 generated feature encoding schemes were entered into t-SNE and shown in **Figure 2**. **Figure 2** clearly depicts that AD and non-AD genes are mixed, indicating that they have limitations to distinguish between AD and non-AD genes in the 2D feature space. It revealed that these features may or may not be able to classify between the AD and non-AD genes. The data pattern also showed that only a widely used linear ML algorithm is not able to classify between the positive and negative samples. Therefore, we have considered all the widely used ML algorithms such as the linear ML classification method with the tree-based method and gradient boosting methods which are known to classify biological problems and other problems were implemented in this study through scikit-learn.

**Figure 2.**
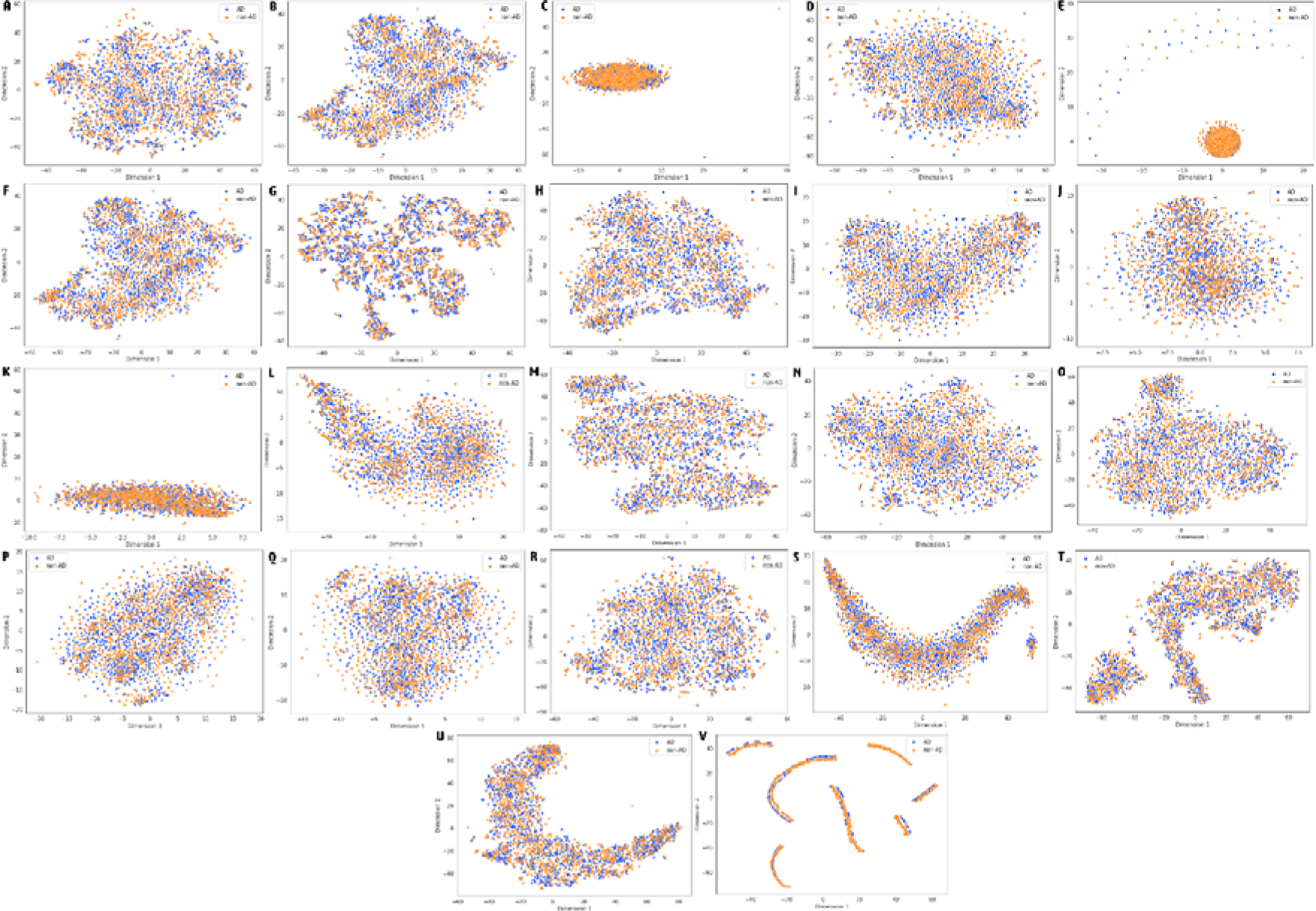
Distribution of the features using t-SNE. (A) AAC, (B) CKSAAP (C) DPC (D) DDE (E) TPC (F) CSKAAGP (G) GAAC (H) GDPC (I) GTPC (J) Geary (K) Moran (L) NMBroto (M) CTDC (N) CTDD (O) CTDT (P) CTriad (Q) KSCTriad (R) QSOrder (S) SOCNumber (T) PAAC (U) APAAC (V) Network features.

### 3.2 Model generation

The performance of 22 different features belonging to eight different groups was assessed by using 16 ML algorithms with 10-fold cross-validation. The accuracy of each feature encoding scheme for different algorithms was recorded and tabulated in **Supplementary Table S2** and **Figure 3**. We have observed that PPI network features achieved excellent performance in several tree and gradient-boosting-based classifiers while sequence-based features are performing worst.

**Figure 3.**
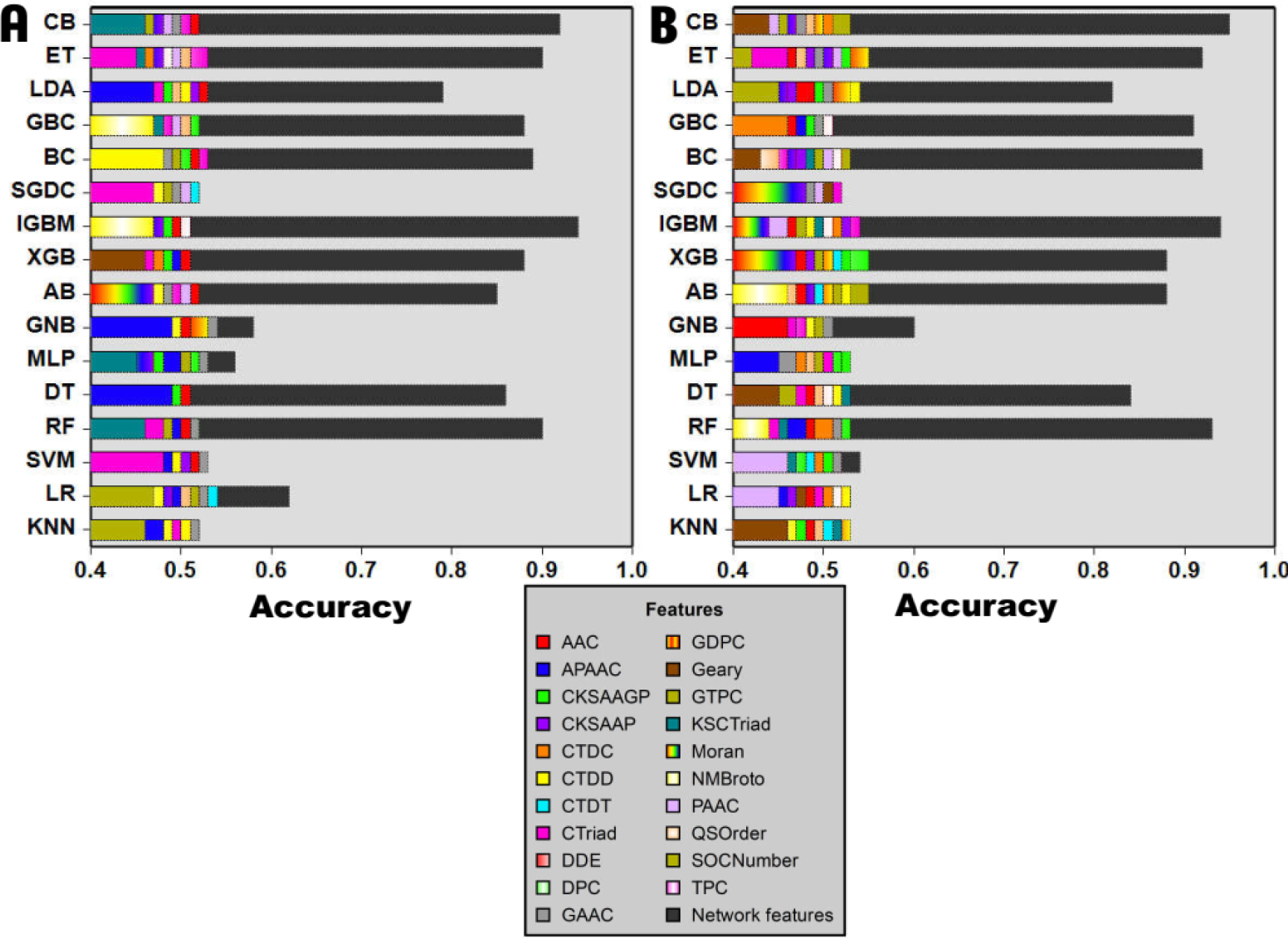
Accuracy of 22 features from 16 ML algorithms. (A) Training accuracy (B) Test accuracy.

We noted that RF, DT, AdaBoost, XGBoost, lightGBM, Bagging Classifier, Gradient Boosting, ExtraTrees, and CatBoost showed accuracy between 85% to 94% for training and test dataset for the network features (**Figure 4**). The sequence-based features showed approximately 50 to 55% accuracy which represents that they are not able to distinguish between the positive and negative datasets. The lightGBM achieved balanced 94% classification accuracy for the training and test dataset respectively for the network-based features. Tree-based methods such as ET and RF both showed 90% accuracy for the training set and 92% and 93% accuracy for the test set respectively. The primary analysis of all the feature encoding schemes revealed that network-based features were performing best while sequence-based features are not able to distinguish between the AD and non-AD genes.

**Figure 4.**
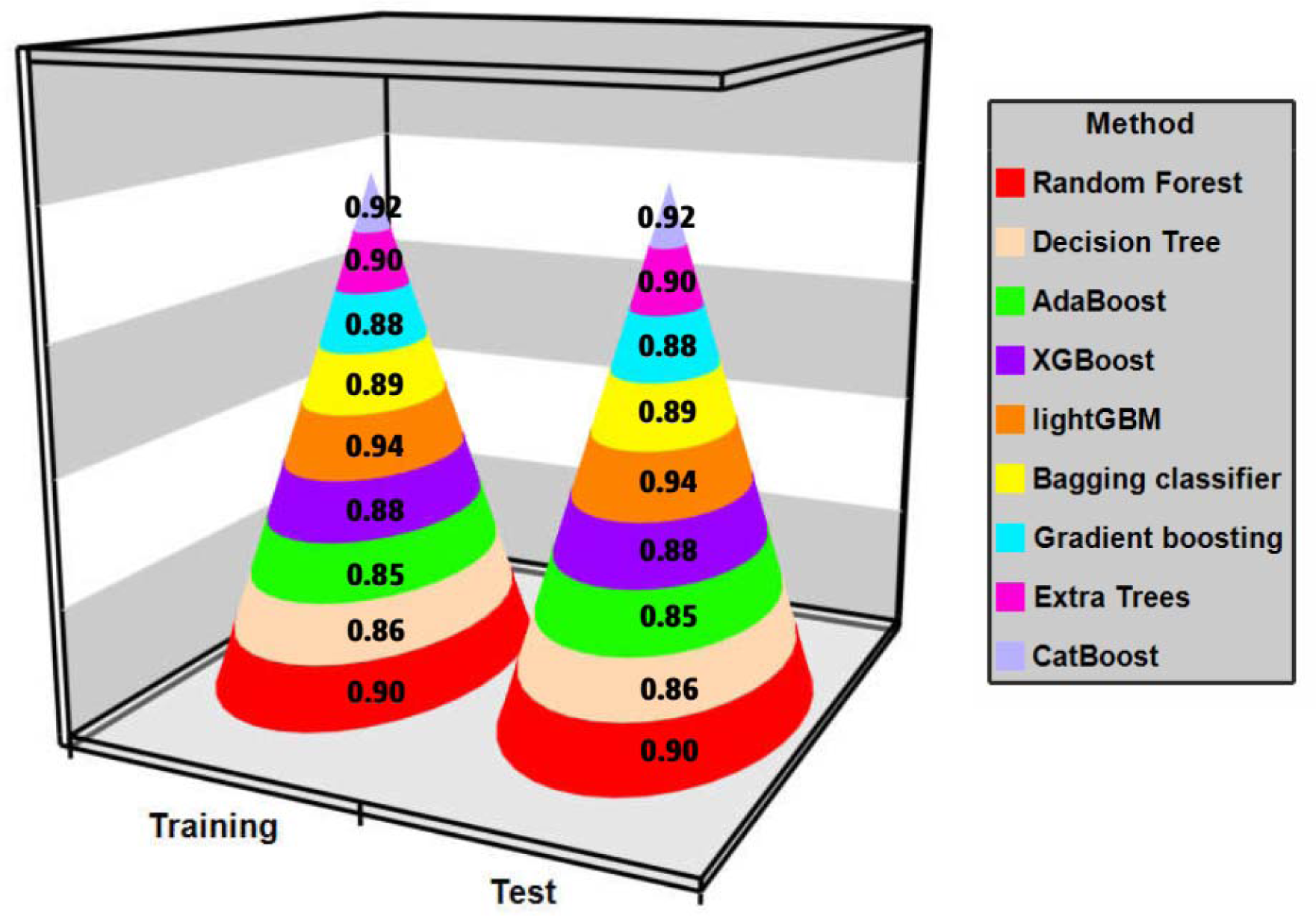
Performance of network-based features using tree-based and ensemble classifier methods.

### 3.3 Optimal feature set selection

The feature selection approach was applied to get the non-redundant and meaningful feature set from various feature encoding schemes. It can reduce the dimension as well as reduce the complexity of the proposed model. The feature selection provides an optimal set of the most important features by using the original dimension of features and can characterize all the relevant information for the respective encoding scheme with fewer but important feature dimensions. A two-step feature selection approach was applied to get the optimal feature set for each feature encoding. Although the sequence-based features were not performing well we have considered those features also for the feature selection process. Because as compared to the PPI network features the calculation of sequence-based features is user-friendly and does not need any third-party software. The network-based features required some database to get their PPI score and then the NetAnalyzer tool to get the topological parameters while sequence-based features can be easily calculated through any programming language by writing the in-house script. The recursive feature selection algorithm wrapped with lightGBM was used and the optimal feature set on the basis of its accuracy score is selected. The name, original feature dimension, the optimal set of features, and lightGBM accuracy were shown in **Table 3**. We observed the accuracy for the selected feature set between 50 to 54.18% for sequence-based features. **Table 3** showed that in sequence-based features the DPC achieved the highest performance of 54.18% while CKSAAP, DDE, and GTPC also achieved 54.00, 54.00, and 53.08% lightGBM feature selection accuracy respectively. In the second step, we assessed the performance of the generated optimal feature set using 16 ML algorithms and observed a 5 to 7% increase in accuracy (**Supplementary Table S3**). The CKSAAP is the only encoding in sequence-based features that showed more than 60% accuracy from CatBoost, lightGBM, LDA, MLP, NB, SVM, and XGBoost methods with overfitting in some methods. After this, the DPC achieved an accuracy of 56.08% and 57.01% for the training and test set respectively from the CatBoost algorithm. The DDE also achieved an accuracy of 57.51 and 55.17% for training and test sets from the Gradient Boosting method. We have seen that feature selection is increasing the accuracy of different feature encoding schemes with less dimension’s datasets (**Supplementary Table S3**). Although this accuracy is not sufficient to construct the ML method based on sequence-based features therefore we moved on to analyzed the network-based feature selection accuracy.

**Table 3.**
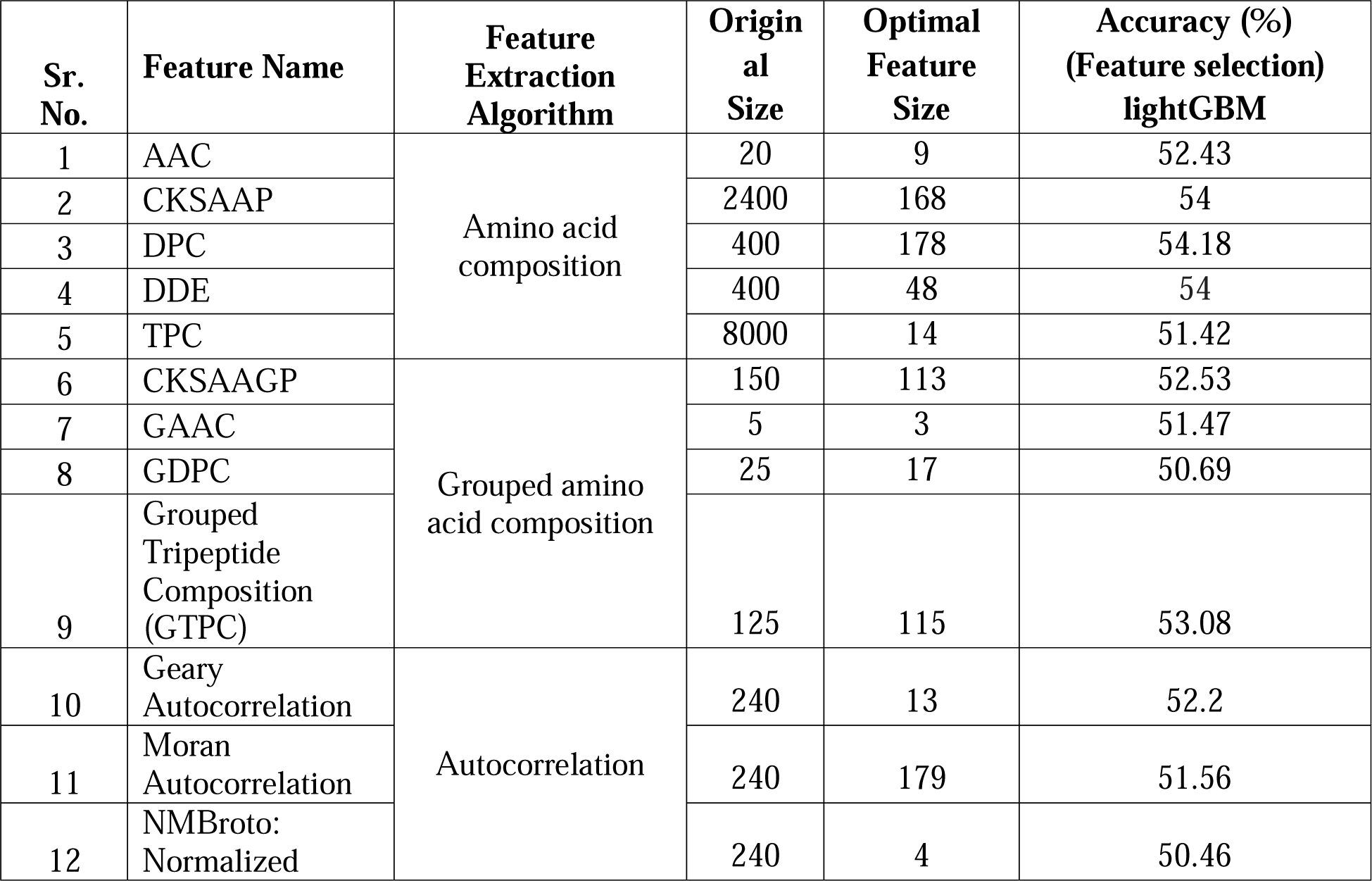
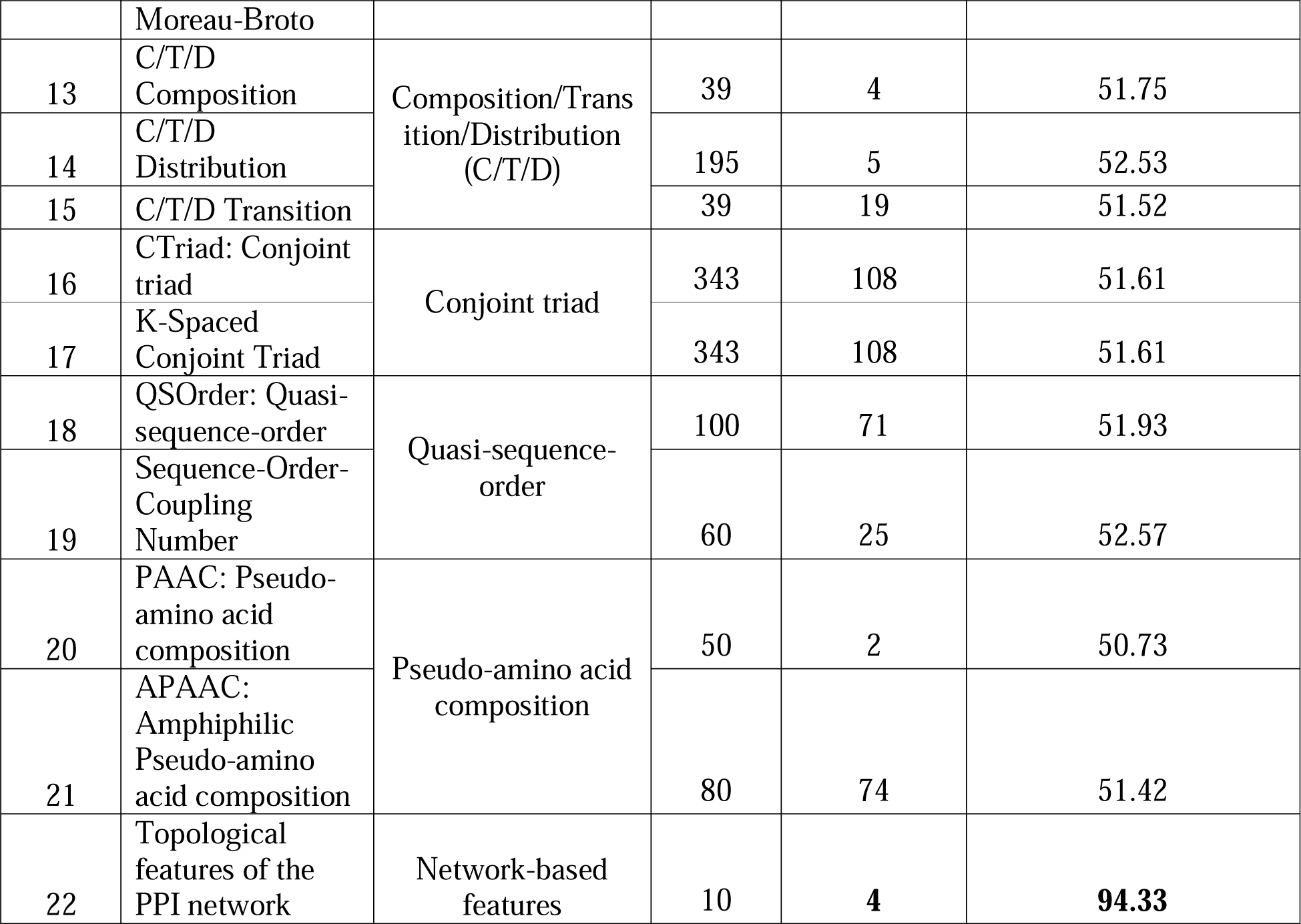
The feature selection accuracy.

The lightGBM produces a 4D feature set out of 10 with 94.33% accuracy. This optimal feature set was employed in 16 ML methods to get a classification model and their accuracy was evaluated. We have seen that feature selection does not increase the accuracy from the original feature set while it decreases the overfitting by reducing the bias between training and test datasets. The accuracy between the training and test set is very balanced in the 4D feature set with a marginal difference as compared to the 10D feature set. The model assessment showed that CatBoost achieved the highest balanced accuracy of 94.87 and 94.94% for training and test datasets respectively. The ExtraTrees and lightGBM also showed approximately 94% accuracy for the training and test sets (**Figure 5**). The results revealed that feature selection is not only significantly improving the accuracy while it is also reducing the overfitting and feature’s dimensions.

**Figure 5.**
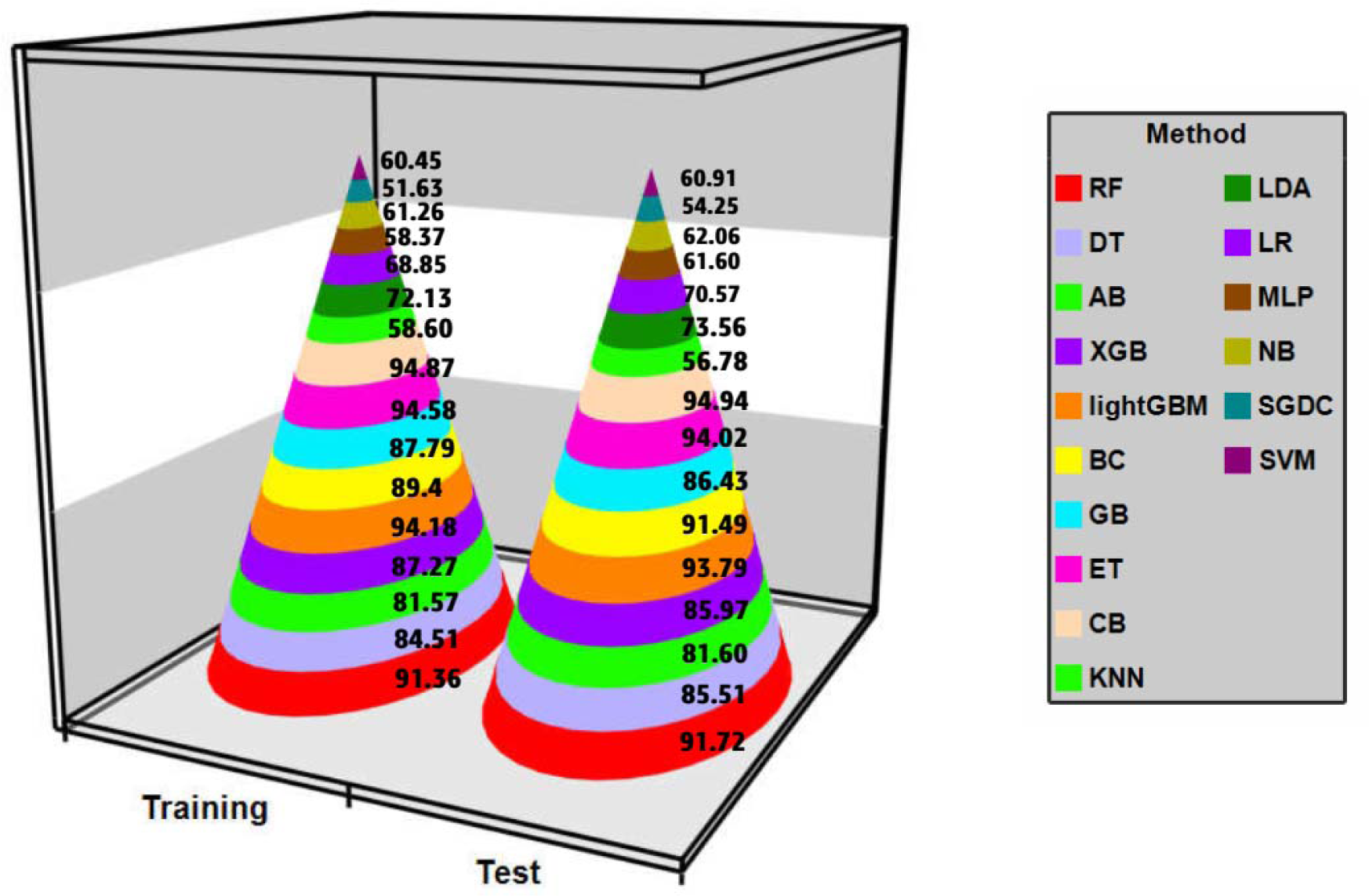
The accuracy (in %) of the 4D feature set from 16 ML methods.

### 3.4 Feature fusion

Feature fusion is a widely used approach to solve the prediction problems demonstrated in several recent studies [62–64]. The network-based features were not fused because they showed ∼94% accuracy with only four features. The two-step feature selection study of the original feature set demonstrated an optimal feature set and its lightGBM prediction accuracy. We have fused only those features which achieved >= 52% accuracy in the first step of feature selection. The CKSAAP, DDE, and DPC showed approximately 54% accuracy in feature selection. The GTPC, Geary, CTDD, and SOC Numbers also showed more than 52% accuracy hence all features were fused to get more accuracy. A total of 24 feature encoding schemes consisting of 6020 features were created from the optimal feature set generated from the original feature set and used for the two-step feature selection process. An optimal feature set was generated in the first step and then fed to 16 ML algorithms for training and test set prediction. The training and test set accuracy with optimal feature dimension is tabulated in **Supplementary Table S4**. We have seen approximately a 6 to 12% increase in the lightGBM feature selection accuracy with approximately 59 to 66% accuracy for 13 feature sets (**Table 4 and Figure 6**).

**Figure 6.**
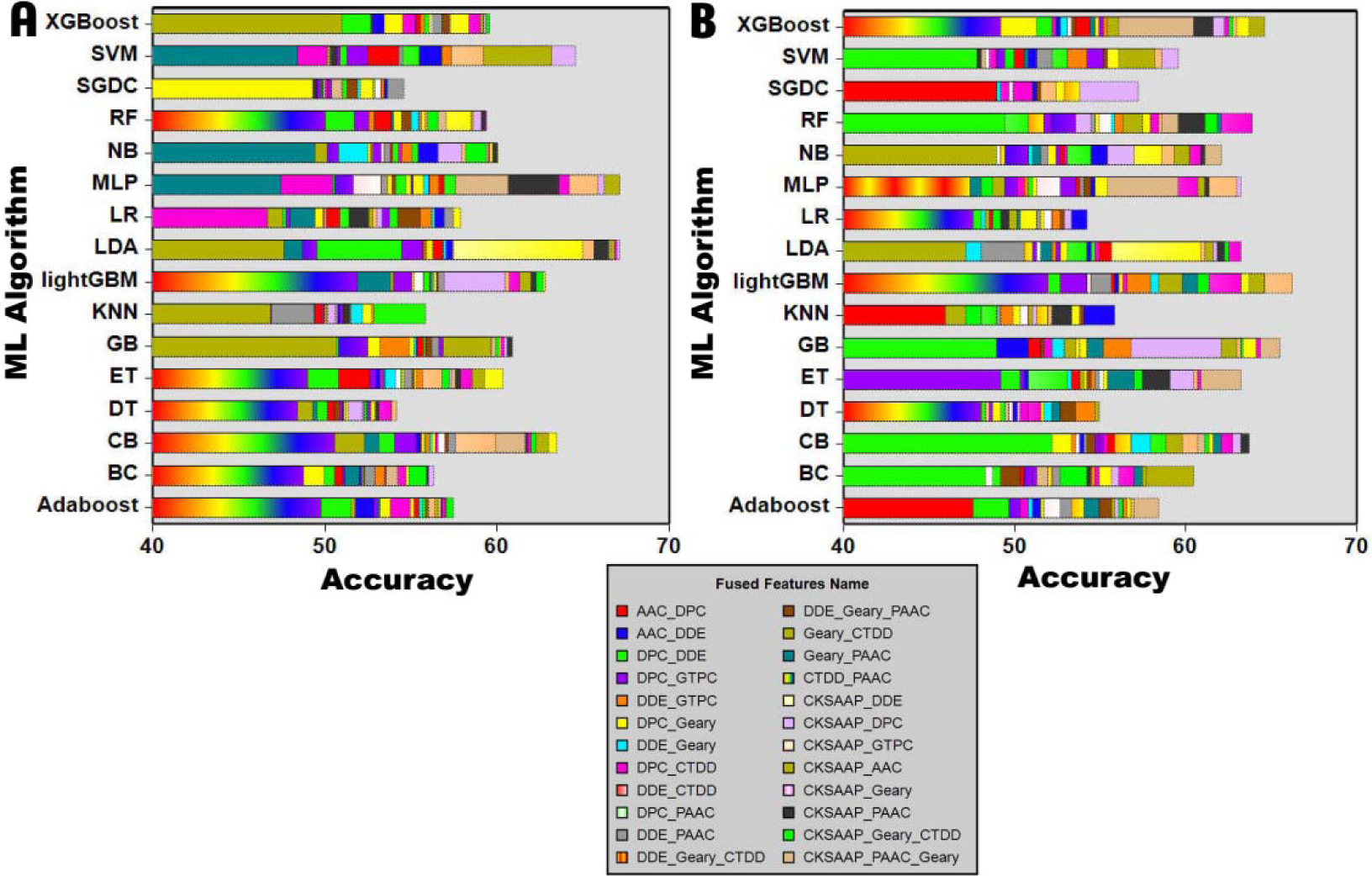
The fused features accuracy was obtained from 16 ML methods. (A) Training data (B) Test data.

**Table 4.** The optimal feature size and lightGBM accuracy for fused features.

| Sr. No. | Feature_Name | Original Size | Optimal Feature size | Accuracy (Feature_selection) |
| --- | --- | --- | --- | --- |
| 1 | AAC_DPC | 187 | 168 | 57.31 |
| 2 | AAC_DDE | 196 | 144 | 56.71 |
| 3 | DPC_DDE | 226 | 182 | 56.99 |
| 4 | DPC_GTPC | 293 | 234 | 56.85 |
| 5 | DDE_GTPC | 163 | 93 | 57.91 |
| 6 | DPC_Geary | 191 | 189 | 57.04 |
| 7 | DDE_Geary | 61 | 60 | 59.11 |
| 8 | DPC_CTDD | 183 | 164 | 57.04 |
| 9 | DDE_CTDD | 53 | 52 | 59.38 |
| 10 | DPC_PAAC | 180 | 151 | 57.54 |
| 11 | DDE_PAAC | 50 | 50 | 59.85 |
| 12 | DDE_Geary_CTDD | 835 | 66 | 59.43 |
| 13 | DDE_Geary_PAAC | 690 | 57 | 59.66 |
| 14 | Geary_CTDD | 435 | 16 | 55.84 |
| 15 | Geary_PAAC | 290 | 13 | 56.76 |
| 16 | CTDD_PAAC | 245 | 7 | 53.45 |
| 17 | CKSAAP_DDE | 216 | 208 | **66.02** |
| 18 | CKSAAP_DPC | 346 | 209 | **63.76** |
| 19 | CKSAAP_GTPC | 283 | 244 | **64.82** |
| 20 | CKSAAP_AAC | 177 | 170 | **65.56** |
| 21 | CKSAAP_Geary | 181 | 180 | **66.06** |
| 22 | CKSAAP_PAAC | 170 | 170 | **66.25** |
| 23 | CKSAAP_Geary_CTDD | 186 | 185 | **65.54** |
| 24 | CKSAAP_PAAC_Geary | 183 | 181 | **64.91** |
|  | **Total** | **6020** | **3193** |  |

We have observed that CKSAAP_PAAC does not lose any dimension in the feature selection and showed the highest lightGBM accuracy (66.25%) as compared to other fused features. All eight features which are fused with CKSAAP are showing more than 60% accuracy (**Accuracy bold in Table 4**). It indicates that the CKSAAP feature can increase accuracy. We have also observed that DDE_PAAC achieved 59.85% on the 50D feature vector. It is the smallest feature dimension as compared to other fused features. Furthermore, the comparison between original and fused feature results from 16 ML algorithms exhibited a significant improvement in accuracy (2-3%). The original features like CKSAAP, DPC, and DDE only got 54% accuracy while the fused features increased the accuracy from 5 to 12% and reached 66% for several feature sets. The accuracy of all the fused features from 16 ML methods were shown in **Figure 6**. Although again this accuracy is not sufficient to propose an ML model. Therefore, to evaluate the performance of these eight fused feature set with the CKSAAP, we have used a hyperparameter tuning approach and generated thousands of ML model, and evaluated them based on the accuracy matrix.

### 3.5 Hyperparameter tuning of sequence-based fused features

Eight fused features were selected with the CKSAAP optimal feature set (n=168) and fed into eight algorithms which showed good accuracy for the fused features. We have tuned multiple parameters (**Supplementary Table S1**) and generated thousands of ML models but we have not found an accuracy >= 70% for any fused features with any method. The accuracy and tuned parameter details for the fused features with the CKSAAP were shown in **Supplementary Table S5**. The training and test accuracy is shown in **Figure 7**. I observed, that the AdaBoost-based method showed the highest accuracy for the test set for CKSAAP_PAAC (69.85%) while for the training set it showed 66.38% accuracy. Although we have not seen more than 70% accuracy for any features, therefore we have not considered and explored the sequence-based features for further analysis.

**Figure 7.**
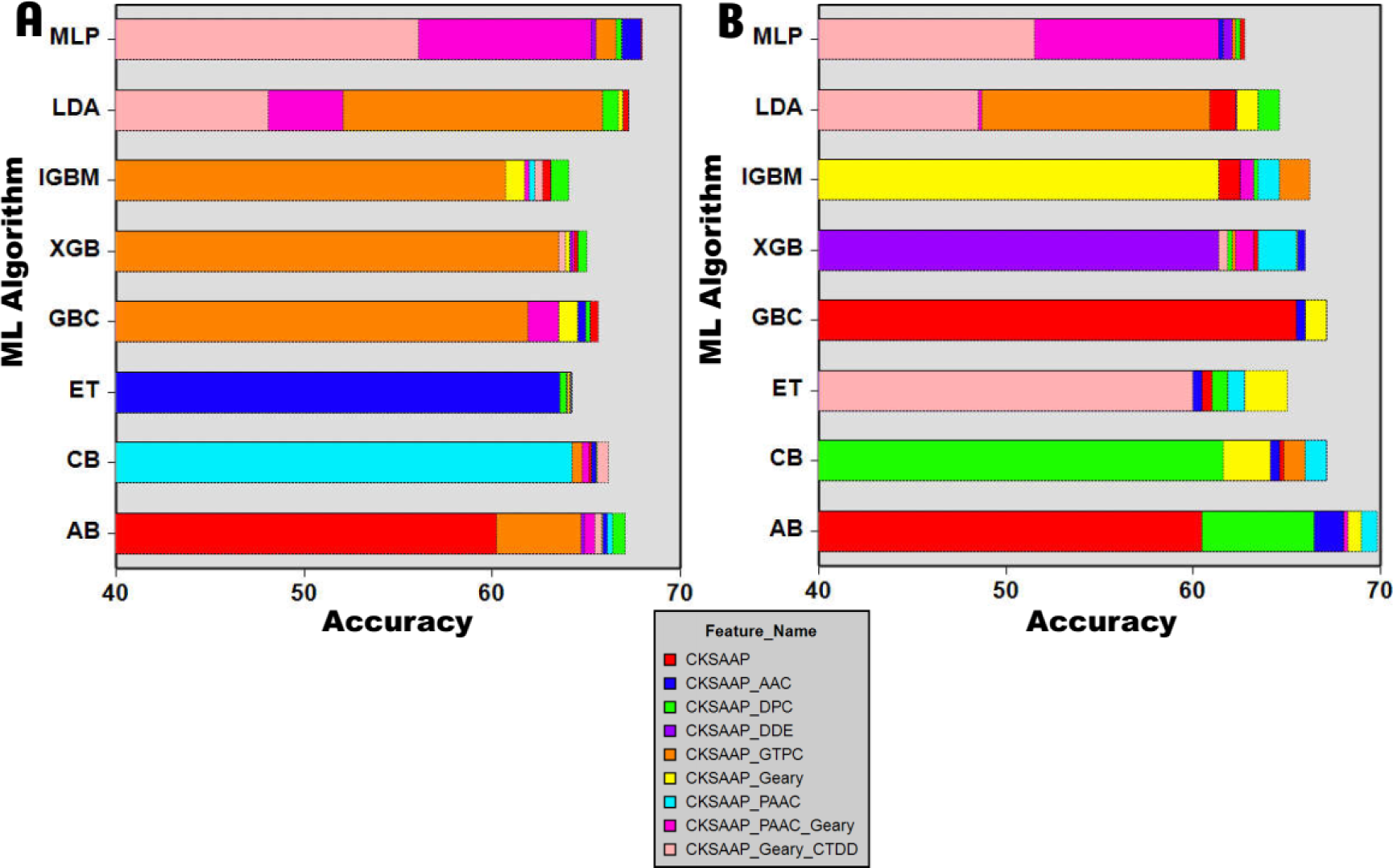
Fused features accuracy (%) obtained by the hyperparameter tuning. (A) Training set (B) Test set.

### 3.6 Network-based feature subset analysis

The above analysis revealed that tree and gradient boosting methods are performing well hence we have considered the best-performing five ML algorithms (RF, lightGBM, XGBoost, ExtraTrees, and CatBoost) for the feature selection and model evaluation on network-based features only. Although the XGBoost is showing less than 90% accuracy, we considered it because, in the initial analysis, it performed once best. We embedded all these algorithms in the wrapper-based feature selection method and used the original network feature set (10D) for feature selection. The 7, 4, 8, 5, and 7 optimal feature sets were identified from RF, lightGBM, XGBoost, ExtraTrees, and CatBoost methods respectively. These optimal feature sets were inputted into these five algorithms with 10-fold CV and their result is reported in **Table 5**. The smallest feature subset with 4 features generated from lightGBM indicated very balanced accuracy for the training and test set. The ET-generated optimal feature subset (5D) also showed good accuracy with training and test set accuracy bias. We have also compared the optimal and original feature set (10D) to assess the performance measurement, whether it is improved or not. The assessment between the optimal and original dataset analysis revealed that optimal features were slightly better performed in RF, ET, and CB classifiers while the lightGBM showed similar performance in the 4D feature set. The XGBoost was performing better in the original dataset while the performance is reduced in the optimal dataset with the 4D feature (**Table 5**). Overall, we have observed that feature selection is improving by 2 to 4% accuracy and reducing the feature dimension also in the training dataset. We identified that only three algorithms (lightGBM, ET, and CB) are showing more than 94% accuracy for the optimal feature set. Therefore, we considered the small feature set consisting of four features and used it for hyperparameter tuning because it is the smallest feature set and also showed very balanced accuracy in training and test datasets (**Table 5**).

**Table 5.**
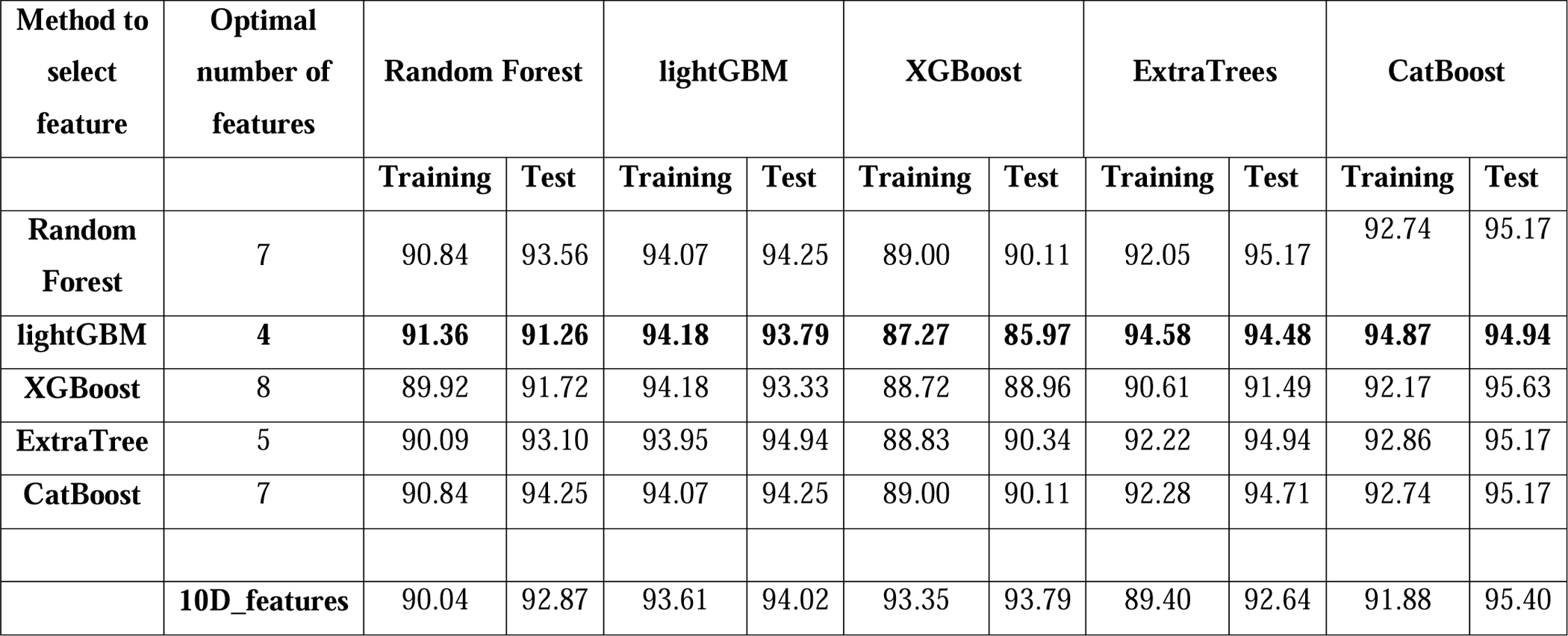
The performance of five ML algorithms on feature selection and their evaluation.

| Method to select feature | Optimal number of features | Random Forest |  | lightGBM |  | XGBoost |  | ExtraTrees |  | CatBoost |  |
| --- | --- | --- | --- | --- | --- | --- | --- | --- | --- | --- | --- |
|  |  | Training | Test | Training | Test | Training | Test | Training | Test | Training | Test |
| Random Forest | 7 | 90.84 | 93.56 | 94.07 | 94.25 | 89.00 | 90.11 | 92.05 | 95.17 | 92.74 | 95.17 |
| lightGBM | 4 | **91.36** | **91.26** | **94.18** | **93.79** | **87.27** | **85.97** | **94.58** | **94.48** | **94.87** | **94.94** |
| XGBoost | 8 | 89.92 | 91.72 | 94.18 | 93.33 | 88.72 | 88.96 | 90.61 | 91.49 | 92.17 | 95.63 |
| ExtraTree | 5 | 90.09 | 93.10 | 93.95 | 94.94 | 88.83 | 90.34 | 92.22 | 94.94 | 92.86 | 95.17 |
| CatBoost | 7 | 90.84 | 94.25 | 94.07 | 94.25 | 89.00 | 90.11 | 92.28 | 94.71 | 92.74 | 95.17 |
|  | **10D_features** | 90.04 | 92.87 | 93.61 | 94.02 | 93.35 | 93.79 | 89.40 | 92.64 | 91.88 | 95.40 |

### 3.7 Optimal model selection

The feature selection results revealed that lightGBM, ET, and CB are performing better than the other two ML algorithms. Therefore, we have considered three algorithms for hyperparameter tuning. They offer many parameters which can be tuned and by tuning those parameters, we have generated thousands of ML models and assessed them for final selection. The tuned parameters are given in **Supplementary Table S1**. The comparison between the previous model and the optimal model revealed that hyperparameter tuning increases the accuracy from 1 to 3% for training and test dataset (**Figure 8 A and B**). We have assessed the optimal models in various statistical parameters using the test dataset and tabulated them in **Table 6**. The CB-based model achieved the best performance with Sens, Spec, Precision, Acc, AUROC, and MCC with their respective values as 96.39, 96.71, 96.83, 96.55, 98.99, and 93.1%. Although the CB model showed slightly less AUROC as compared to other algorithms, however, it outperformed in all other measured parameters (**Figure 9**). The overall result revealed that CB based model is better than lightGBM and ET. It indicates that optimal model selection was a better approach than other approaches, therefore the CB-based model was further selected for implementation. The 4D network-based features are provided in **Supplementary Table S6** for the reconstruction of the proposed method.

**Figure 8.**
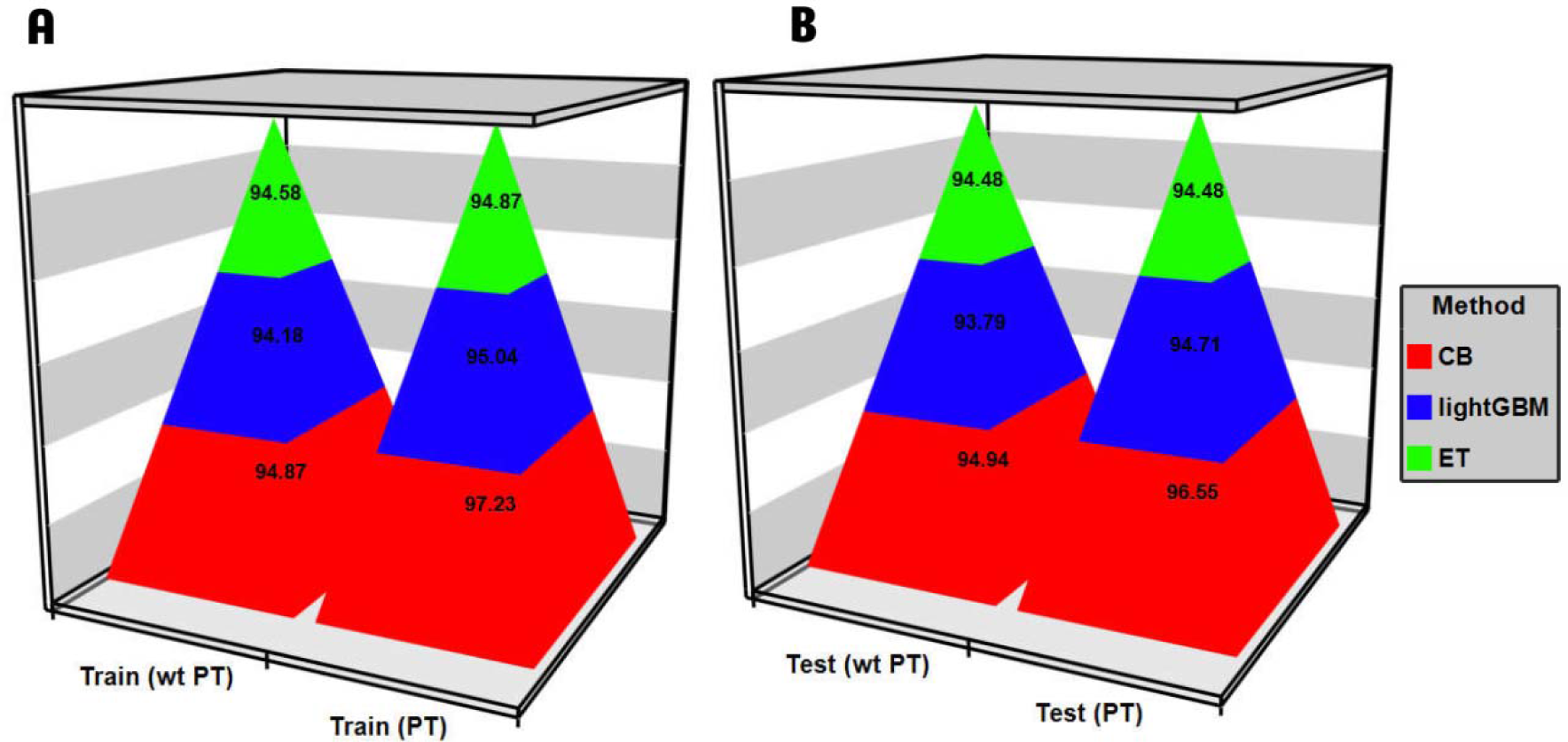
Accuracy comparison between tuned models *vs.* non-tuned modes. (A) training (B) test.

**Figure 9.**
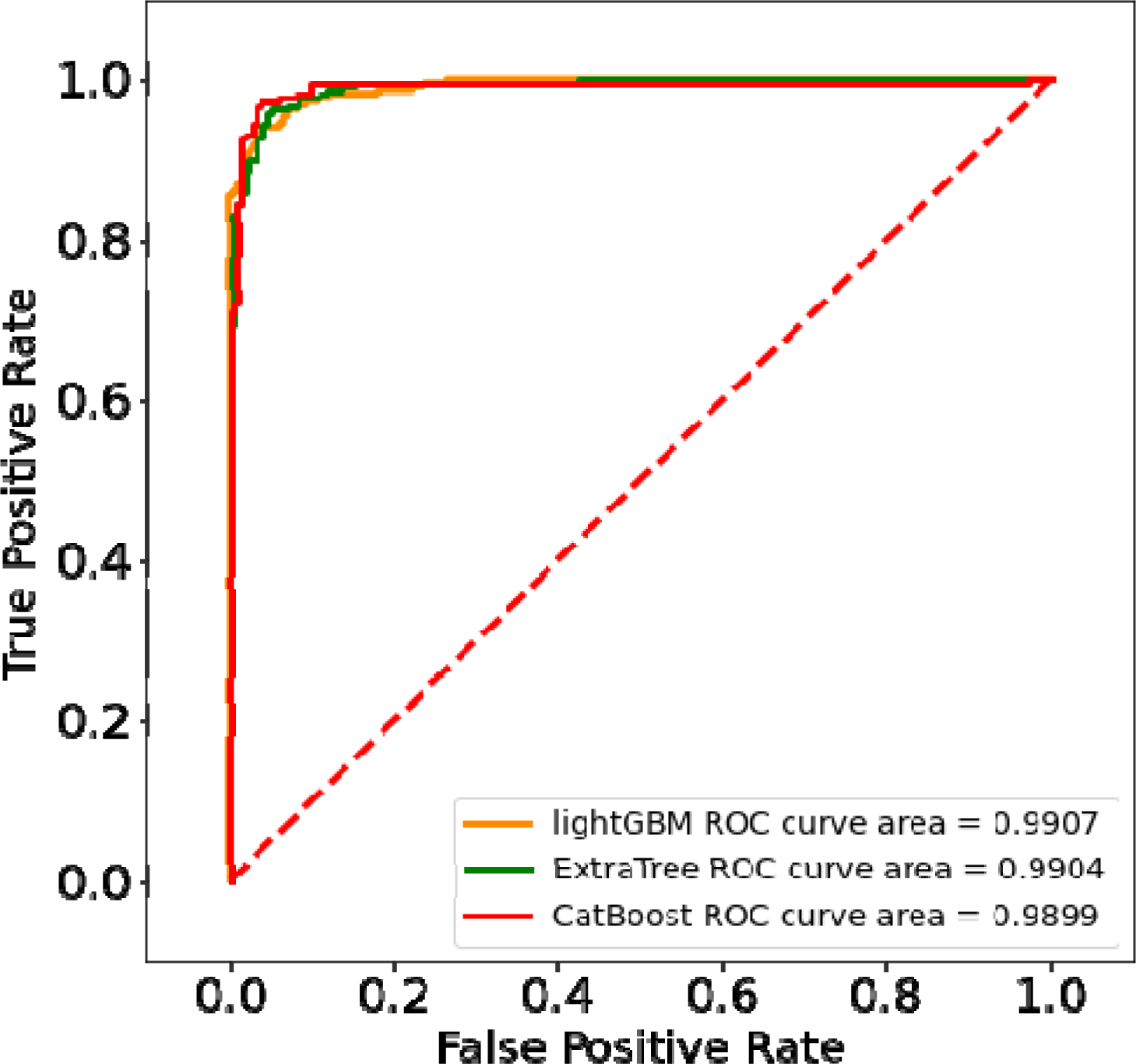
ROC curve for three final methods selected after rigorous exercise of feature selection hyperparameter tuning and model generations.

**Table 6.** The performance of three selected machine learning techniques on the network-based features. l2_lr = l2_leaf_reg, LR = learning rate, mf = max_features, msl = min_samples_leaf, ss = min_samples_split and ne = n_estimators.

| Model | Hyperparameters | Sens | Spec | Precision | Acc | AUROC | MCC |
| --- | --- | --- | --- | --- | --- | --- | --- |
| CB | depth = 8, iteration = 250,<br>l2_lr = 3, LR = 0.3 | 96.39 | 96.71 | 96.83 | 96.55 | 98.99 | 93.1 |
| lightGBM | depth = 3, iteration = 150,<br>l2_lr = 3, LR = 0.2 | 93.24 | 96.24 | 96.27 | 94.71 | 99.07 | 89.47 |
| ET | mf = 3, msl = 2, mss = 3,<br>ne = 526 | 93.69 | 95.3 | 95.41 | 94.48 | 99.04 | 88.98 |

### 3.8 Evaluation of AlzGenPred using independent datasets

During the model training, over-fitting is also possible to achieve higher accuracy. Hence it is highly recommended that the proposed model should be evaluated on the unseen dataset that whether it is capable of accurately classifying the AD and non-AD genes. Notably, the blind dataset should not be used during the training of the model (model development). Hence, we have ensured that the independent dataset is not present in the training procedure and the sequences which were present during training were removed from the independent dataset.

#### 3.8.1 AlzGene Database

The AlzGene dataset consists of the experimentally identified AD genes generated from systematic meta-analyses [53]. The database has 695 genes with their score. We retrieved those genes and mapped them to the Uniprot to get the sequence. We fetched 642 sequences and the sequence which were present in the training or test dataset were removed. Finally, 502 sequences were selected and inputted into the STRING database and then PPI is retrieved. Then the network parameters of 477 genes were used as input to the AlzGenPred. Our proposed method outperforms the blind dataset with 96.43% accuracy. It correctly classified 460 AD genes while incorrectly classified only 17 genes. It indicates that our proposed method can classify AD and non-AD genes (**Figure 10A**).

**Figure 10.**
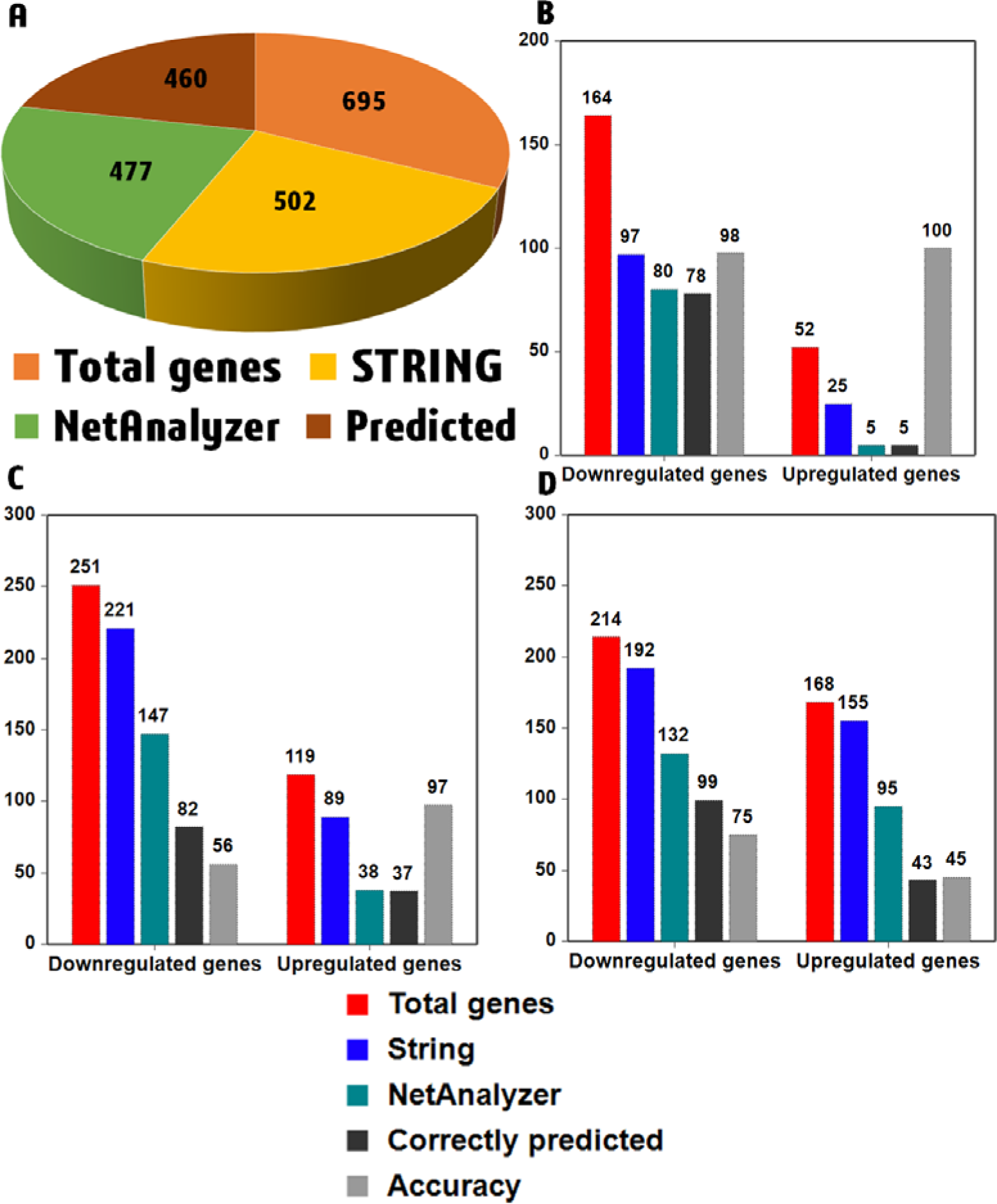
Validation of the AlzGenPred tool from different independent datasets. (A) AlzGene dataset (B) GSE113437 (C) GSE67333 and (D) GSE162873.

#### 3.8.2 Evaluation of AlzGenPred using GEO datasets

We have downloaded and pre-processed the RNA sequencing dataset generated from AD and healthy controls and their upregulated and downregulated genes were inputted into the developed method for the server validation.

##### 3.8.2.1 GSE113437

The data is pre-processed using the steps described in the material and method section and then the read counts per million, PCA, and heatmaps were generated from the normalized count data (**Supplementary Figure 1**). The upregulated (52) and downregulated genes (164) were identified (**Supplementary Figure 2A**). MA plot and the enriched pathways for the DEGs were identified (**Supplementary Figure 2B and C**). Out of 52 upregulated genes, only 25 genes were mapped to the STRING database. We have downloaded the PPI map and inputted it into the NetAnalyzer plugin for feature calculation. It kept only five genes and predicted the five genes features. These five genes were inputted to the AlzGenPred server where we found 100% prediction accuracy (**Supplementary Figure 3A**). After that, 164 downregulated genes were inputted into the STRING database and then it mapped 97 genes. Out of 97 genes, the features of 80 genes were predicted of which 78 were correctly classified by our proposed method with 97.5% accuracy (**Supplementary Figure 3B**). The detail of the genes and prediction accuracy is shown in **Figure 10B**. We have also carried out the gene set enrichment analysis where we predicted the pathways for the downregulated genes and upregulated genes. A tree of gene set enrichment analysis consisting of pathways information for up and down-regulated genes was also identified and shown in **Supplementary Figure 3C**. It represents that identified genes are involved in various pathways which actively participate in AD progression.

##### 3.8.2.2 GSE67333

The data is pre-processed and the read counts per million, PCA, and heatmaps were generated and shown in **Supplementary Figure 4**. After that, we predicted the upregulated and downregulated genes (**Supplementary Figure 5A**). The MA plot and the enriched pathways for the DEGs were identified (**Supplementary Figure 5B and C**). These upregulated genes (119) were inputted into the STRING database and from there we found 89 genes PPI network. That PPI network was inputted into the NetAnalyzer plugin from where we obtained the features for 38 genes. Out of the 38 genes, the proposed method can correctly classify 37 genes with 97.36% accuracy (**Supplementary Figure 6A**). Then the downregulated genes (251) were inputted into the STRING database and from there we obtained 221 genes PPI network. That PPI network was inputted into the NetAnalyzer plugin from where we obtained the features for 147 genes. Out of the 147 genes, our proposed method correctly classified 82 genes with 55.76% accuracy (**Supplementary Figure 6B**). The detail of the genes and prediction accuracy is shown in **Figure 10C**. The gene set enrichment analysis revealed that the up and down-regulated genes are involved in the key pathways which play a key role in AD progression (**Supplementary Figure 6C**).

##### 3.8.2.3 GSE162873

The data is pre-processed and then the read counts per million, PCA, and heatmap were generated and shown in **Supplementary Figure 7**. Then we calculated the upregulated and downregulated genes (**Supplementary Figure 8A**) with the MA plot (**Supplementary Figure 8B and C**). Here we have taken a total of eight samples for AD (four) and healthy control (four) generated from two ES-derived neural cells from two AD patients. Then the common up (168) and downregulated (214) genes were taken from both samples and fed to the STRING database (**Supplementary Figure 8D**). It mapped 155 upregulated genes and 192 downregulated genes. These genes were inputted into the NetAnalyzer plugin where we have seen that 95 and 132 genes features were predicted. Out of 95 genes, only 43 genes were correctly classified with 45.26% (**Supplementary Figure 9A**) while for downregulated genes we have seen that 99 genes were correctly classified out of 132 genes with 75% accuracy (**Supplementary Figure 9B**). The detail of the genes and prediction accuracy is shown in **Figure 10D**. Then we also predicted the pathways using the gene set enrichment analysis where we have seen that the predicted genes are expressed in the key pathways (**Supplementary Figure 9C**).

### 3.9 Standalone tool

We have also developed a standalone tool for users with a well-documented tutorial for the users. A snapshot of the tool is shown in **Figure 11**. The user can download the tool from https://www.bioinfoindia.org/alzgenpred/ and https://github.com/shuklarohit815/AlzGenPred.

**Figure 11.**
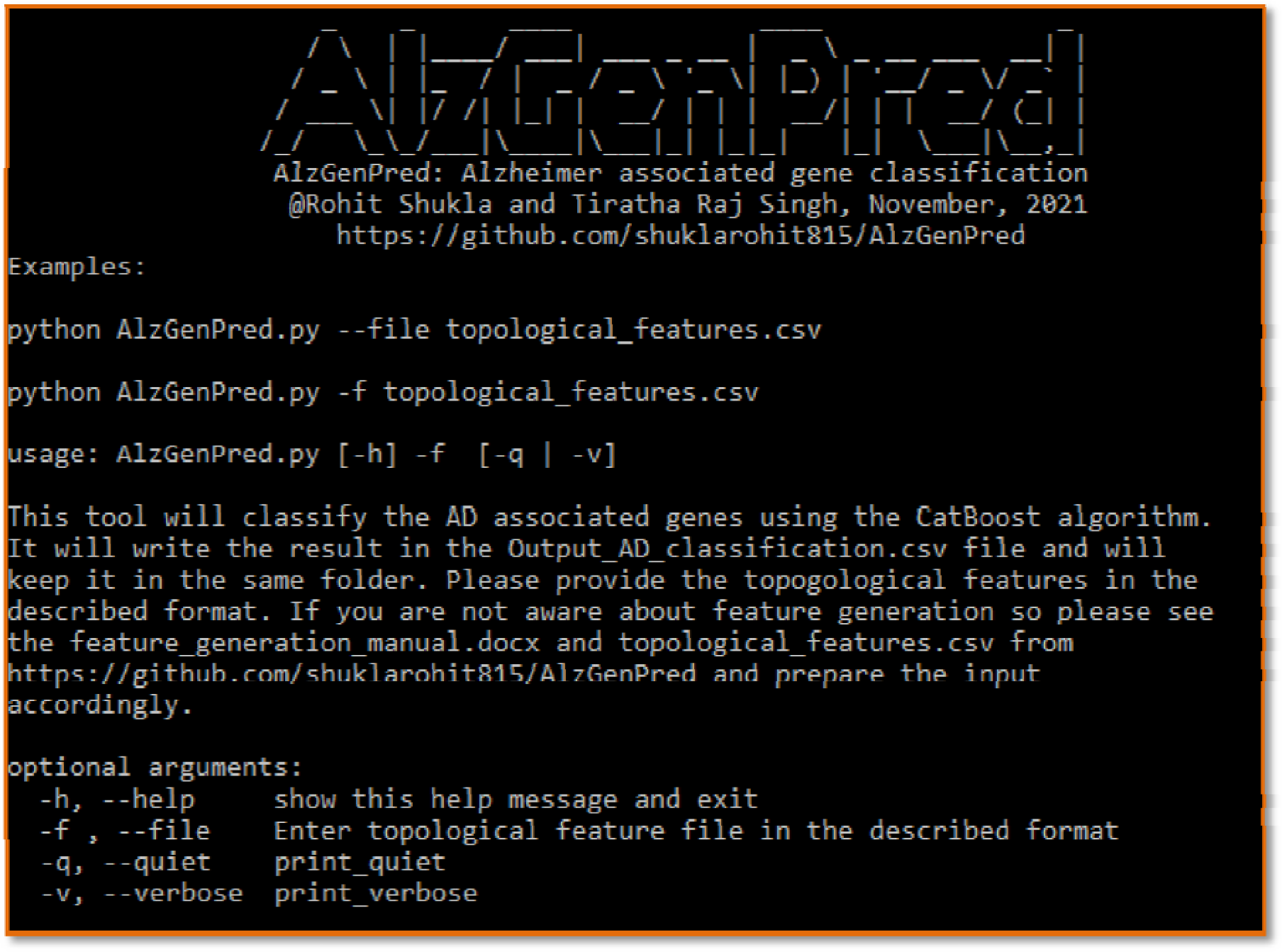
The GUI of the AlzGenPred tool with detailed help.

## 4. Discussion

AD is a common cause of dementia and impaired cognitive function. It is still an incurable disease despite the advancement of science. Therefore, scientists are continuously looking to identify potential biomarkers and drug targets for AD. It is a most challenging neurodegenerative disease due to its complicated and progressive mechanism. There are very few genes information available, which does not cover a significant portion of human genes. AD can be caused by several risk factors and several genes are involved in AD progression. There is a lot of efforts are necessary to identify the new genes for AD which can act as a biomarker for the diagnosis of the disease. Machine learning is a newly emerging technique, which can learn and identify the pattern from complex data and can identify the potential genes which are involved in several diseases. In the current study, we have used the ML approach and developed a standalone tool that can identify AD-associated genes. We have retrieved the AD-associated genes dataset from the DisGeNET and then pre-processed it. The pre-processing is a very important step that can remove the overfitting of the machine learning model [19,62]. Then 1086 genes were used as a training and test set. By using all those genes, we have calculated 13,504 features consisting of sequence and network-based features. We have utilized the widely used 16 machine learning algorithms and faded the calculated features into it. Here, we have seen that network-based features can classify AD-associated genes while sequence-based features are not able to classify the AD-associated genes. In a previous study also the network-based features can classify the AD genes [19]. Then we used the feature selection and hyperparameter tuning approach against the selected features. The previous studies uses feature selection and got good accuracy for the ML models [51,65]. Despite all these approaches, the sequence-based features were not able to classify the AD and non-AD genes. Hence, we focused on the network-based features and developed a standalone tool for the AD *vs.* non-AD gene classification with 96.55% accuracy for the test dataset. Model validation is a critical process in proposing a new method [66,67]. Hence, we have also taken the real dataset from the AlzGene database [53] and validated our model. It showed 96.43% accuracy for the AlzGene dataset. Then we also validated our model with the transcriptomics dataset where also we have seen that the proposed method can identify the new AD genes from the large dataset. An AD classification method was proposed by Jamal *et al.* which is also using the network features while AlzGenPred showed more accuracy than the existing method [19]. A sequence-based machine learning model was also proposed which can classify the AD-associated genes but they have not proposed any pipeline to classify the user-given genes [68]. Here also AlzGenPred outperformed with good accuracy. We have proposed the method to the scientific community https://www.bioinfoindia.org/alzgenpred/ and https://github.com/shuklarohit815/AlzGenPred as a standalone tool, which was not provided by the previous studies. In this study, we tried to develop an ML-based method that can classify the AD-associated genes and also proposed its standalone version so the user can download this method and can classify the AD genes from the large dataset.

## 5. Conclusion

AD is a neurological disorder that cannot be identified in its early stage. An enormous amount of multi-omics data is available which contains thousands of AD-associated genes but the participation of these genes in AD is still not deciphered. Therefore, we have developed a CatBoost-based method to classify the AD *vs.* non-AD genes. We have taken 13,504 features and inputted them into 16 ML algorithms which reveal that network-based features can classify between the AD and non-AD genes while sequence-based features are not able to distinguish between the AD and non-AD genes. The lightGBM-based feature selection method was also applied to reduce the feature dimensions. Then the feature fusion approach was used and 24 different features were constructed however we have not seen any significant improvement in the accuracy. Even hyperparameter tuning of the sequence-based fused features did not increase the accuracy by more than 70%. Therefore, we have proposed a CatBoost-based method using network-based features called AlzGenPred. It got 97.02% and 96.55% accuracy for the training and test dataset. After that, we also validated the proposed method using the independent dataset and transcriptomics dataset. The validation in the blind dataset revealed that the proposed method can correctly identify the biomarkers which can be further used for the *in-vitro* and *in-vivo* validation.

## Declaration of Competing interest

The authors declare that there are no competing interests.

## Author’s contributions

TRS conceived the study. RS carried out all the experiments and the data analysis. RS and TRS participated in the overall design and coordination of the study. The first draft of the manuscript was prepared by RS. Both authors read and approved the final manuscript.

## Acknowledgment

RS and TRS want to thank the ICMR (ISRM/11(53)/2019) for providing the Senior Research Fellowship to RS.

## Notes

### Competing Interest Statement

The authors have declared no competing interest.

