## Supplementary_Figures for "*AlzGenPred*: A CatBoost based method using network features to classify the Alzheimer’s Disease associated genes from the high throughput sequencing data"

**Running title**

Classification of potential Alzheimer’s Disease genes

Rohit Shukla ^1^**^#^** and Tiratha Raj Singh ^1,2^*

^1^Department of Biotechnology and Bioinformatics, Jaypee University of Information Technology (JUIT), Waknaghat, Solan, H.P., 173234, India

^2^Centre for Excellence in Healthcare technologies and Informatics (CEHTI), Jaypee University of Information Technology (JUIT), Waknaghat, Solan, H.P., 173234, India

**^#^**Current Address: Center of Excellence for Aging and Brain Repair, University of South Florida, Morsani College of Medicine, Tampa, FL, USA

**^*^Corresponding author**

Dr. Tiratha Raj Singh,

Associate Professor,

Department of Biotechnology and Bioinformatics,

Jaypee University of Information Technology,

Waknaghat, Solan, 173215, H.P., India

**Keywords:** Alzheimer’s disease, Neurofibrillary tangles, Machine learning, CatBoost, Network features.

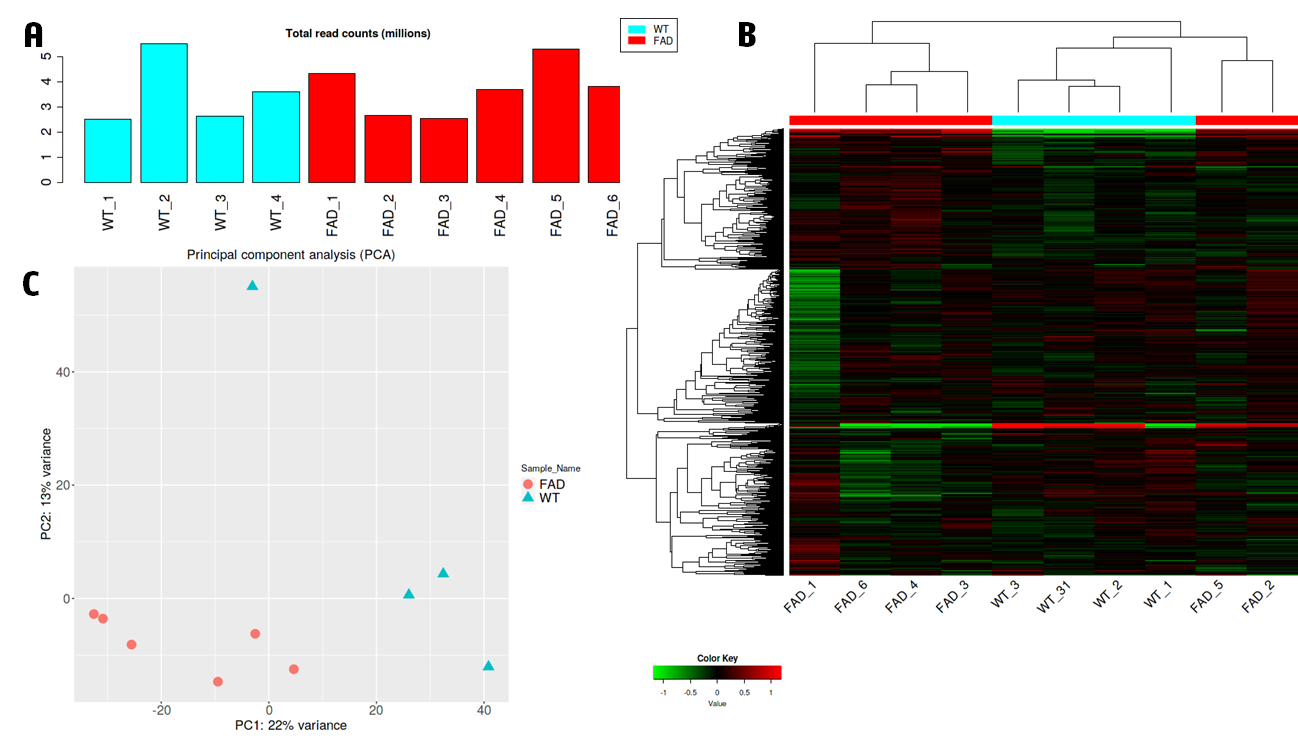

**Supplementary Figure 1:** Quality control parameters of RNAseq data. (A) Total read counts in million (B) Heatmap for the AD *vs.* healthy controls (C) Principal component analysis.

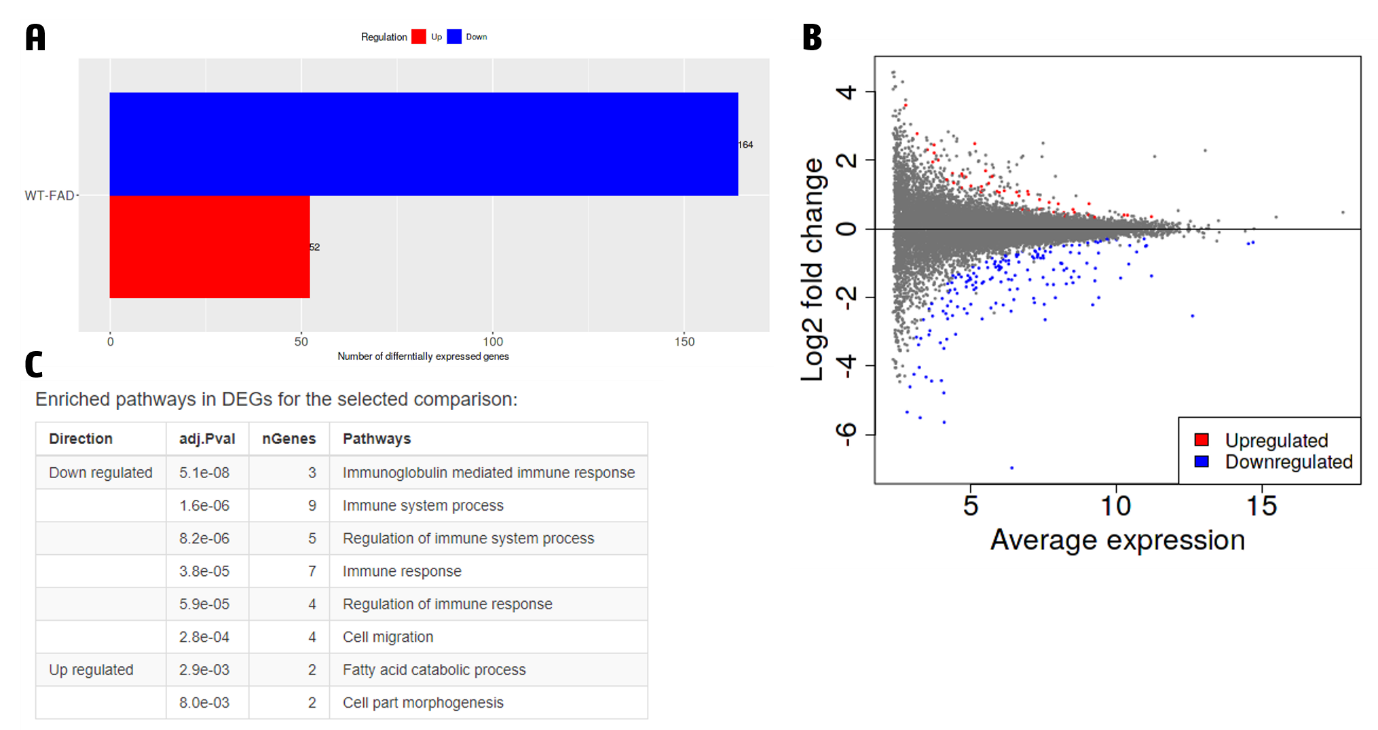

**Supplementary Figure 2:** (A) Up and downregulated genes (B) MA plot for differentially expressed genes (C) Pathway enrichment analysis.

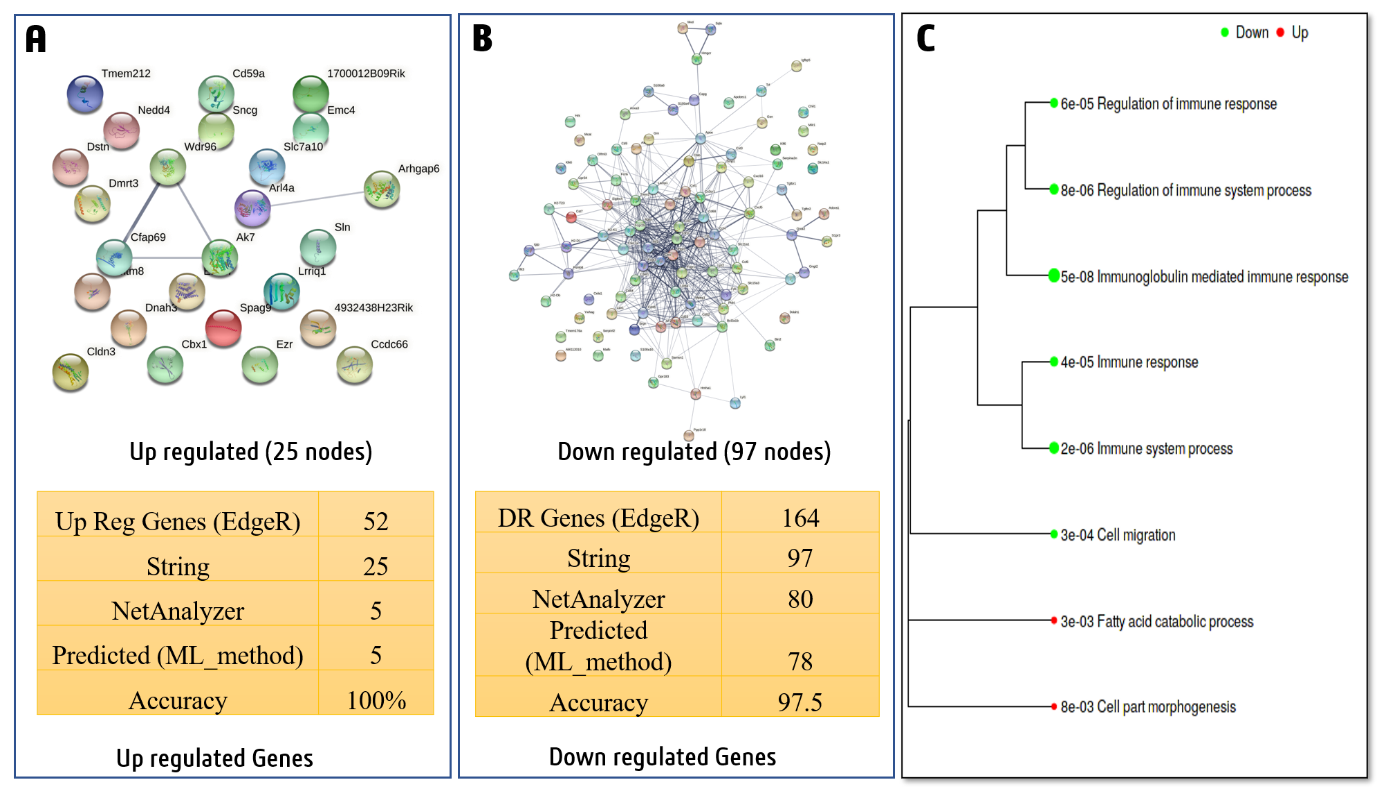

**Supplementary Figure 3:** Classification of the up and down regulated genes using AlzGenPred. (A) Up regulated genes (B) Down regulated genes (C) Pathway analysis for up and down regulated genes.

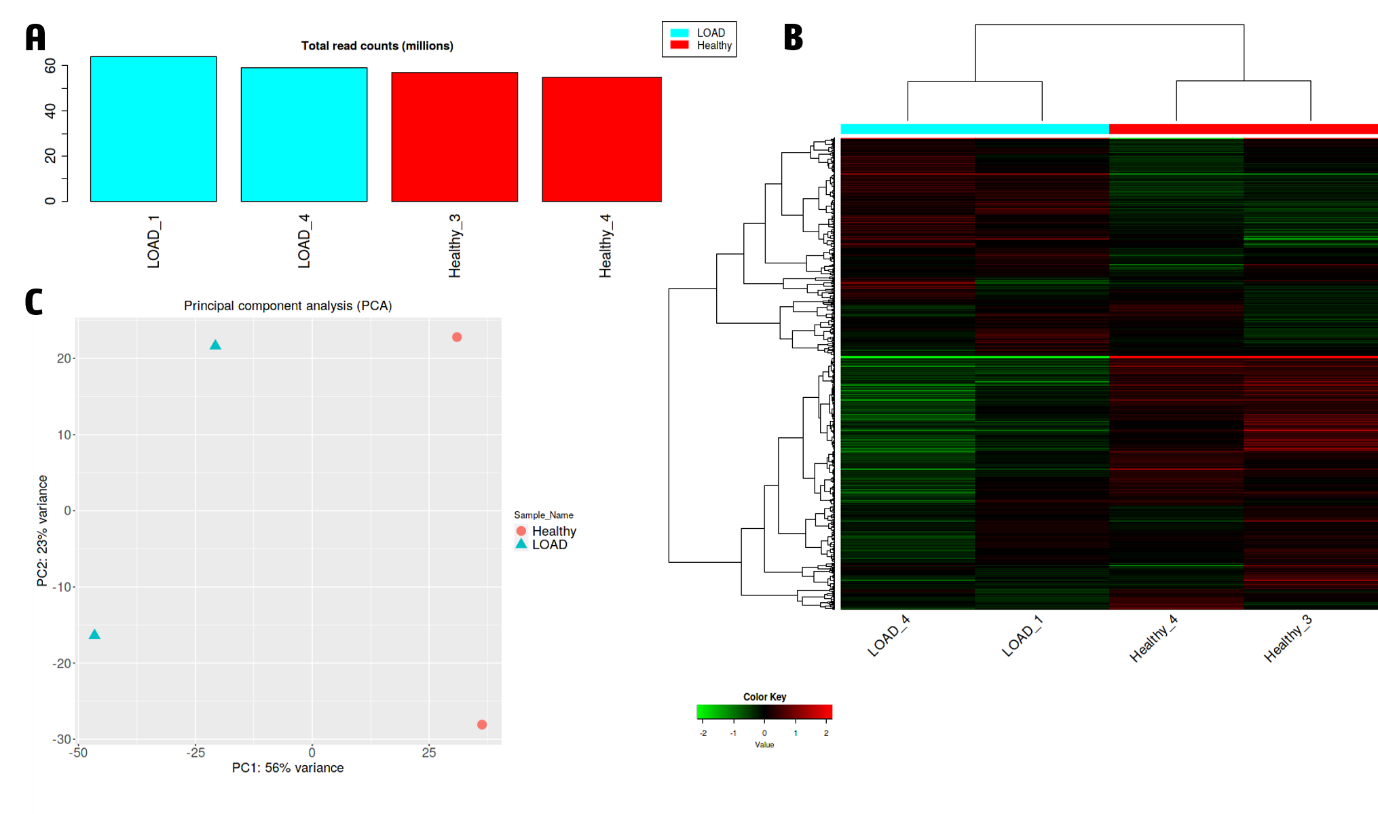

**Supplementary Figure 4:** Quality control parameters of RNAseq data. (A) Total read counts in million (B) Heatmap for the AD *vs.* healthy controls (C) Principal component analysis.

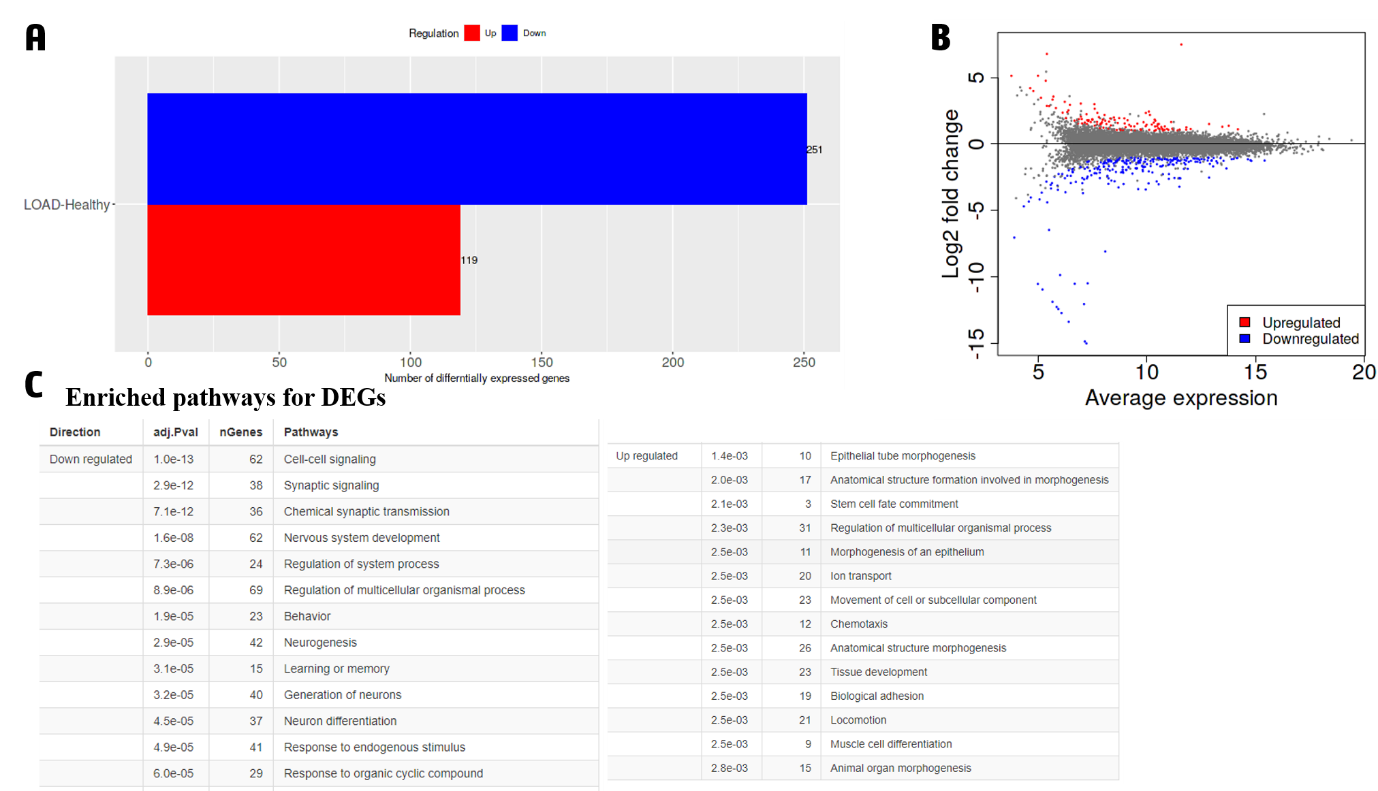

**Supplementary Figure 5:** (A) Up and down regulated genes (B) MA plot for differentially expressed genes (C) Pathway enrichment analysis.

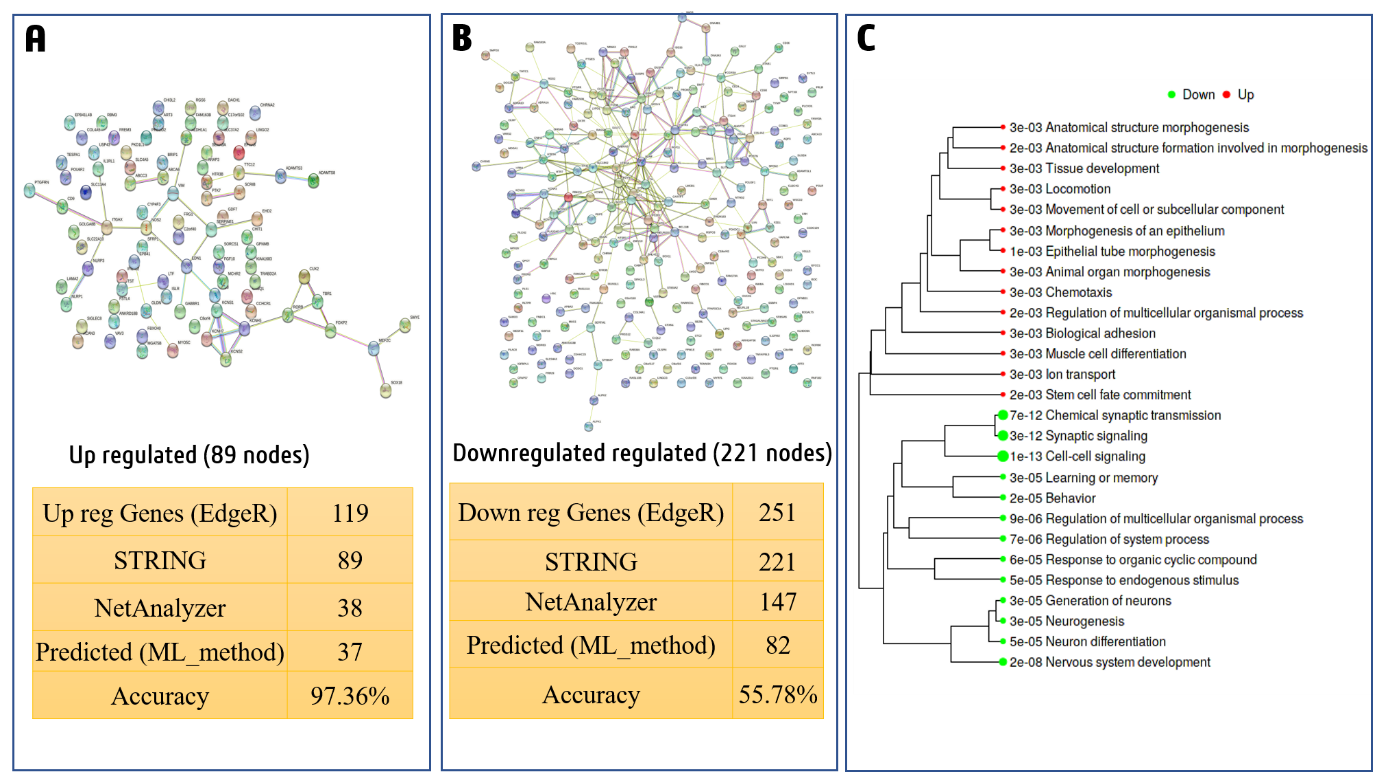

**Supplementary Figure 6:** Classification of the up and down regulated genes using AlzGenPred. (A) Up regulated genes (B) Down regulated genes (C) Pathway analysis for up and down regulated genes.

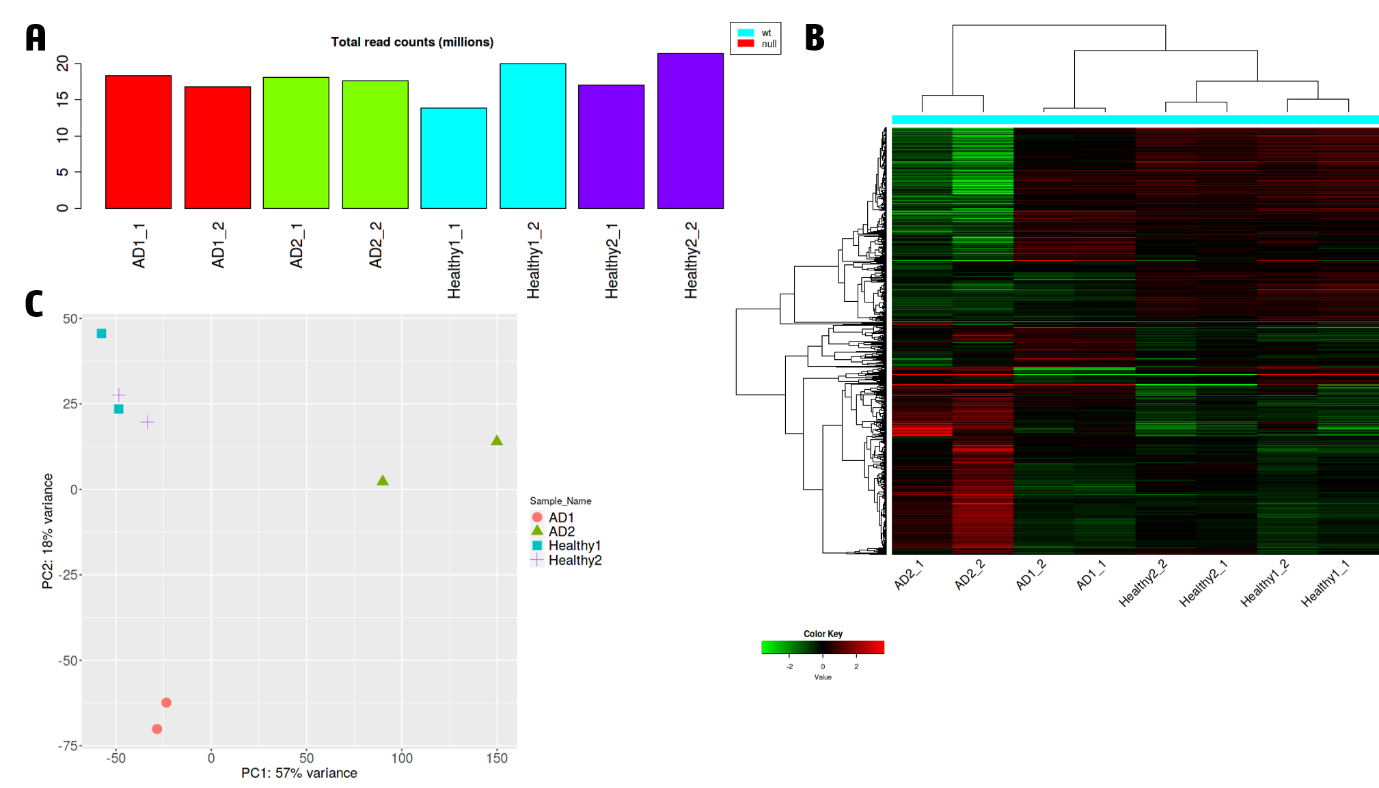

**Supplementary Figure 7:** Quality control parameters of RNAseq data. (A) Total read counts in million (B) Heatmap for the AD *vs.* healthy controls (C) Principal component analysis.

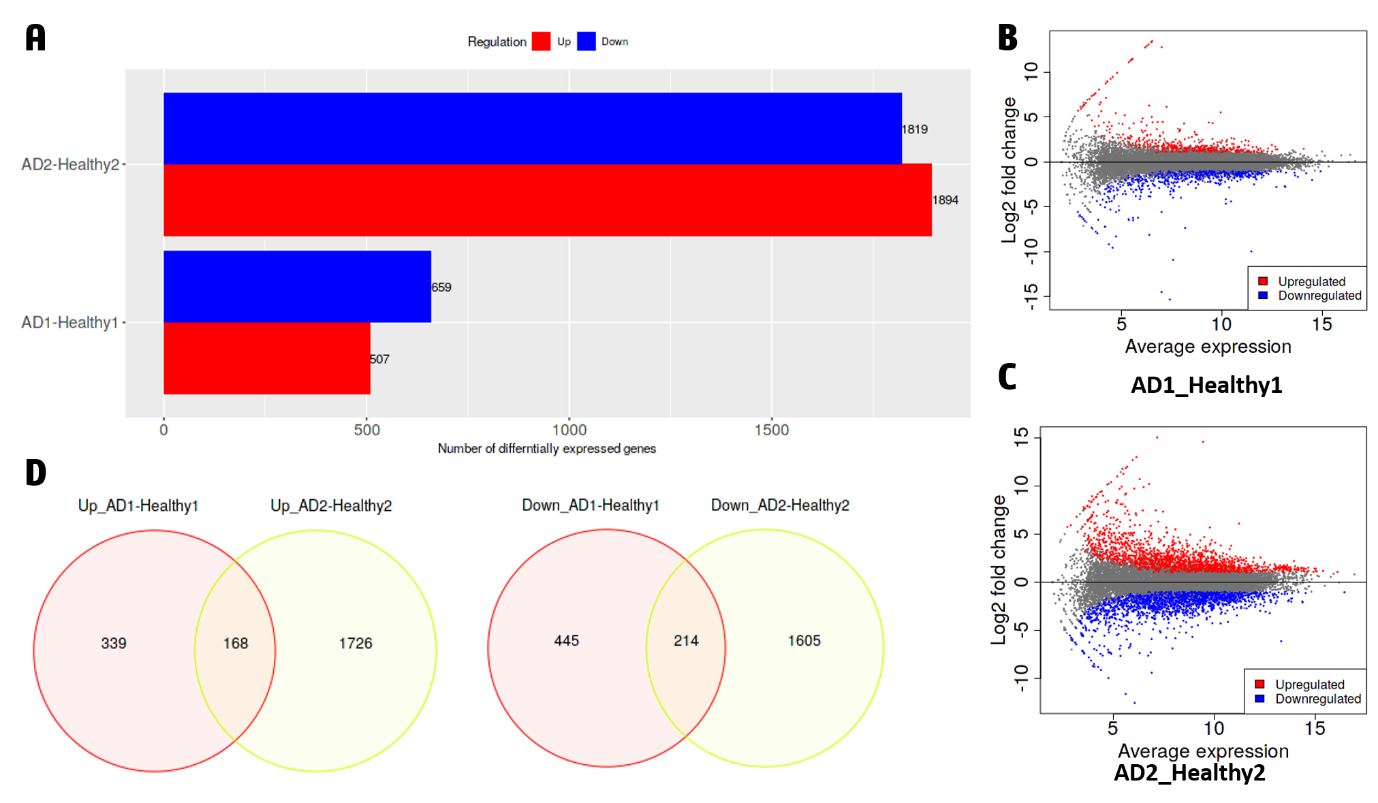

**Supplementary Figure 8:** (A) Up and downregulated genes (B) MA plot for differentially expressed genes for first two samples (C) MA plot for differentially expressed genes for second two samples (D) Venn diagram represent the common genes between 2 AD and 2 healthy controls.

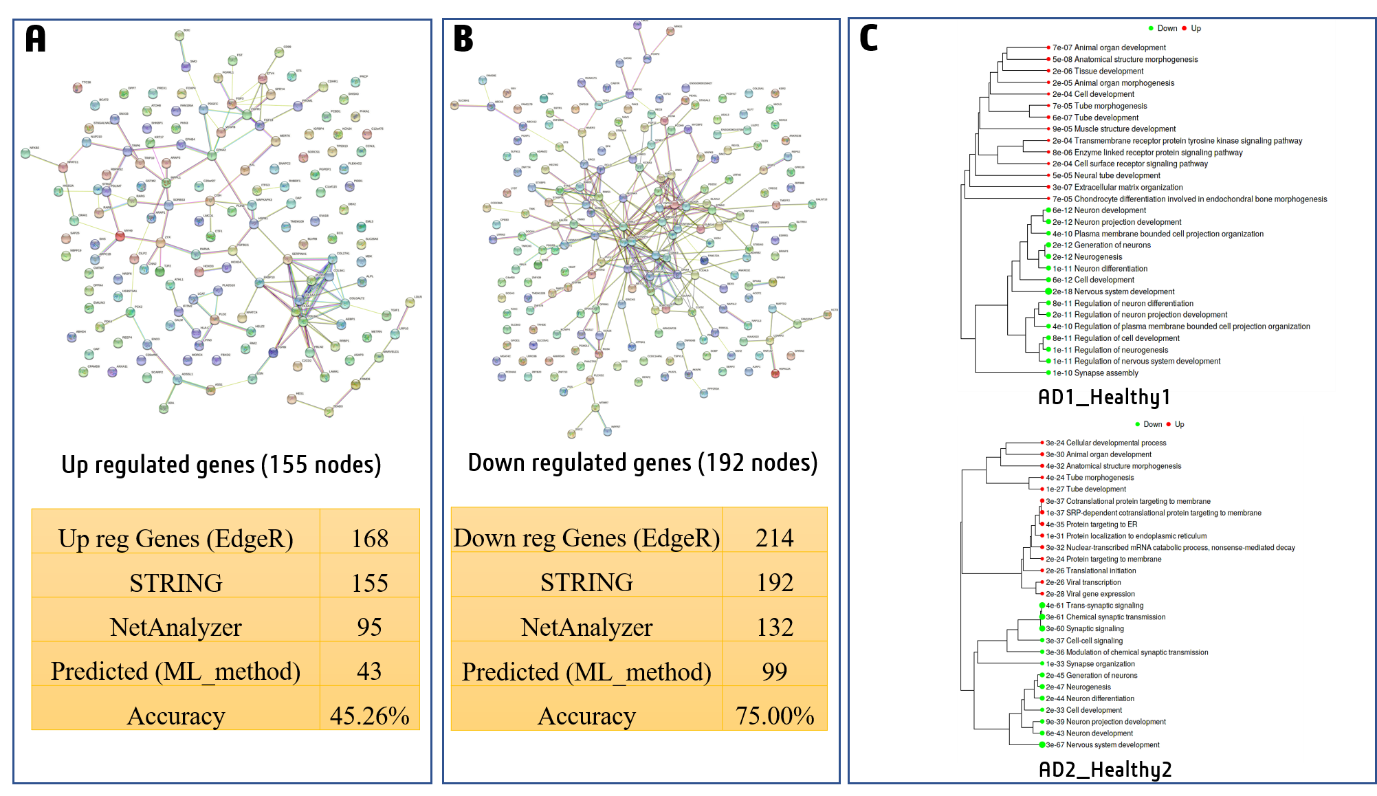

**Supplementary Figure 9:** Classification of the up and down regulated genes using AlzGenPred. (A) Up regulated genes (B) Down regulated genes (C) Pathway analysis for up and down regulated genes.
